# Cooperative and antagonistic interactions between sub-clones favour the co-existence of multiple resistance mechanisms in melanoma

**DOI:** 10.1101/2025.10.23.684170

**Authors:** Karin Schlegelmilch, Viola Hollek, Steven Hooper, Giovanni Giangreco, Sasha Bailey, Sarah Macfarlane, Alexandrine Carminati, Louis Tully, Amy Bowes, Stephanie Strohbuecker, Ben Shum, Samra Turajlic, Xiao Fu, Erik Sahai

## Abstract

Intra-tumour heterogeneity is a major obstacle to durable responses to targeted cancer therapy, yet how different resistant cell states interact within the same tumour remains poorly understood. In this study, we demonstrate cooperativity between co-occurring resistant states in a single tumour. Using BRAF mutant melanoma as a paradigm, we generate three different resistant states within a single model and demonstrate that they exhibit varying differentiation states and migratory capacities and share few common therapeutic vulnerabilities. Through a combination of experiments, including using Cre-mediated recombination to generate heterogeneity in existing tumours, and *in silico* modelling, we show that intra-tumour heterogeneity is the most favoured state for therapy resistant tumours. This is underpinned by signalling between different melanoma states, with YAP1 active cells providing supporting signals for other cells but inhibiting their own proliferation. Optimal disease control requires targeting both the YAP1 active cell state and the inter- cellular communication networks. We identify the histone demethylase inhibitor GSK-J4 as being particularly effective in targeting both features of resistant tumours and demonstrate its ability to control melanoma with multiple concurrent resistance mechanisms.

## Introduction

Therapies targeting driver oncogenes, such as BRAF in melanoma and EGFR in lung cancer, achieve striking initial responses but rarely deliver durable disease control (Long *et al*., 2017; Sosman *et al*., 2012; Kobayashi *et al*., 2005). Numerous mechanisms for resistance to targeted therapy have been proposed. These include the existence of rare pre-existing subclones with mutations that confer resistance, selection for resistant mutants that arise during therapy, adaptation of signalling networks, supporting signals from the tumour microenvironment, and transitions in cell state (Marine, Dawson and Dawson, 2020; Ramos and Bentires-Alj, 2015). In BRAF-mutant cutaneous melanoma, mutations that lead to reactivation of ERK signalling, such as those in NRAS or MEK, are observed in 50% of cases (Moriceau *et al*., 2015; Shi *et al*., 2014). Non-genetic mechanisms involving transcriptional activation of YAP1 and transitions to both melanocytic and mesenchymal cell states can lead to resistance to combined BRAF and MEK inhibition (Lin *et al*., 2015; Verfaillie *et al*., 2015; Arozarena and Wellbrock, 2019; Marine, Dawson and Dawson, 2020; Rambow, Marine and Goding, 2019). Signals emanating from the tumour microenvironment, notably the extracellular matrix (ECM), also provide protection from targeted therapy. It is likely that all of these mechanisms play a role in certain contexts (Fedorenko *et al*., 2016; Titz *et al*., 2016; Marusak *et al*., 2020; Hirata *et al*., 2015). Many studies have identified vulnerabilities of drug resistant states, including targeting of receptor tyrosine kinases, cell adhesion signalling, YAP1, histone deacetylases, and redox state (Labrie *et al*., 2022; Sun *et al*., 2014; Wang *et al*., 2018; Sharma *et al*., 2010; Goyal *et al*., 2023; Kim *et al*., 2016; Boshuizen *et al*., 2018; Hugo *et al*., 2015). However, it is not clear if these strategies are equally effective against different resistance mechanisms. In addition, intra-tumour heterogeneity can hinder effective immune responses against tumours (Wolf *et al*., 2019).

Recent work has highlighted that multiple tumour cell transcriptional and epigenetic states can co-exist, in both treatment naïve and treated disease (Marine, Dawson and Dawson, 2020). For example, melanoma cells can be found in neural crest, mesenchymal, and melanocytic states within the same tumour (Rambow *et al*., 2018). Moreover, clinical evidence indicates that multiple genetic resistance mechanisms are frequently present upon disease recurrence in a patient (Spain *et al*., 2023). Thus, a ‘one size fits all’ approach to tackling therapy resistance is unlikely to be effective and it is now imperative to devise strategies that are effective against multiple mechanisms of resistance that may co-exist in the same patient. Critically, cell plasticity means that targeting one resistant state can increase the prevalence of others, making it essential to understand how different resistant states interact (Rambow *et al*., 2018). Despite this, the majority of pre-clinical studies have focused on individual mechanisms of resistance in isolation, with little comparative analysis across different modes of resistance and no systematic exploration of how diverse resistant states cooperate or compete within the same tumour.

In this study, we establish three different mechanisms of BRAF and MEK inhibitor resistance within a single isogenic model of human melanoma, enabling direct comparison their drug sensitivity, both in terms of acquired vulnerability and synergistic or additive combinations. We proceed to establish a model in which the different modes of resistance can be generated within an established tumour. This reveals bi-directional communication between cells with high and low YAP1 activity. Combined experimental and computational modelling establishes that YAP1 high cells are ‘super-competitors’ but also provide paracrine support signals to YAP1 low cells. As a result, the emergence of parallel mechanisms of resistance is favoured. We identify that the optimal treatment regime for these heterogeneous tumours is targeting of both the YAP1 high state and the paracrine signals that support YAP1 low cells, and demonstrate that this can be achieved with the histone demethylase inhibitor GSK-J4. Together, these findings provide a mechanistic explanation for the heterogeneity in cell state observed in therapy resistant disease and identify an actionable strategy to overcome it in malignant melanoma.

## Results

### Heterogeneity in cell state is a feature of therapy resistant melanoma

To explore intra-tumour heterogeneity in therapy resistant tumours, we stained sub- cutaneous lesions from patients with BRAF mutant melanoma who received combinations of BRAF and MEK inhibitors and immunotherapy with markers of cell state (cohort from PEACE study detailed in (Spain *et al*., 2023)). These included indicators of melanoma differentiation (PMEL, MLANA, CDH1 and SOX10), proliferation (Ki67), amoeboid state (PDPN) (de Winde *et al*., 2021), modulators of immunotherapy response (HLA-ABC and PD- L1), and cell adhesion and signalling molecules previously implicated in therapy resistance (FN1, TNC, YAP1 (Hirata *et al*., 2015; Diazzi, Tartare-Deckert and Deckert, 2023; Lin *et al*., 2015)). Even within small regions of individual tumours, we observed considerable heterogeneity in the expression of these markers (Supp. Fig. 1A). Principal component analysis indicated that these cells reflected a continuum, as previous proposed (Arozarena and Wellbrock, 2019; Rambow, Marine and Goding, 2019), and that nuclear YAP1 contributed positively to the first principal component (Supp. Fig. 1B). In addition to this local heterogeneity, prior genomic analysis indicated that multiple genetic events linked to resistance also occurred within the same patient (Spain *et al*., 2023).These data demonstrate that heterogeneity is a feature of therapy resistant tumours but cannot discriminate whether intra-tumour heterogeneity is actively maintained, or merely a transient consequence of ongoing tumour evolution.

**Figure 1.**
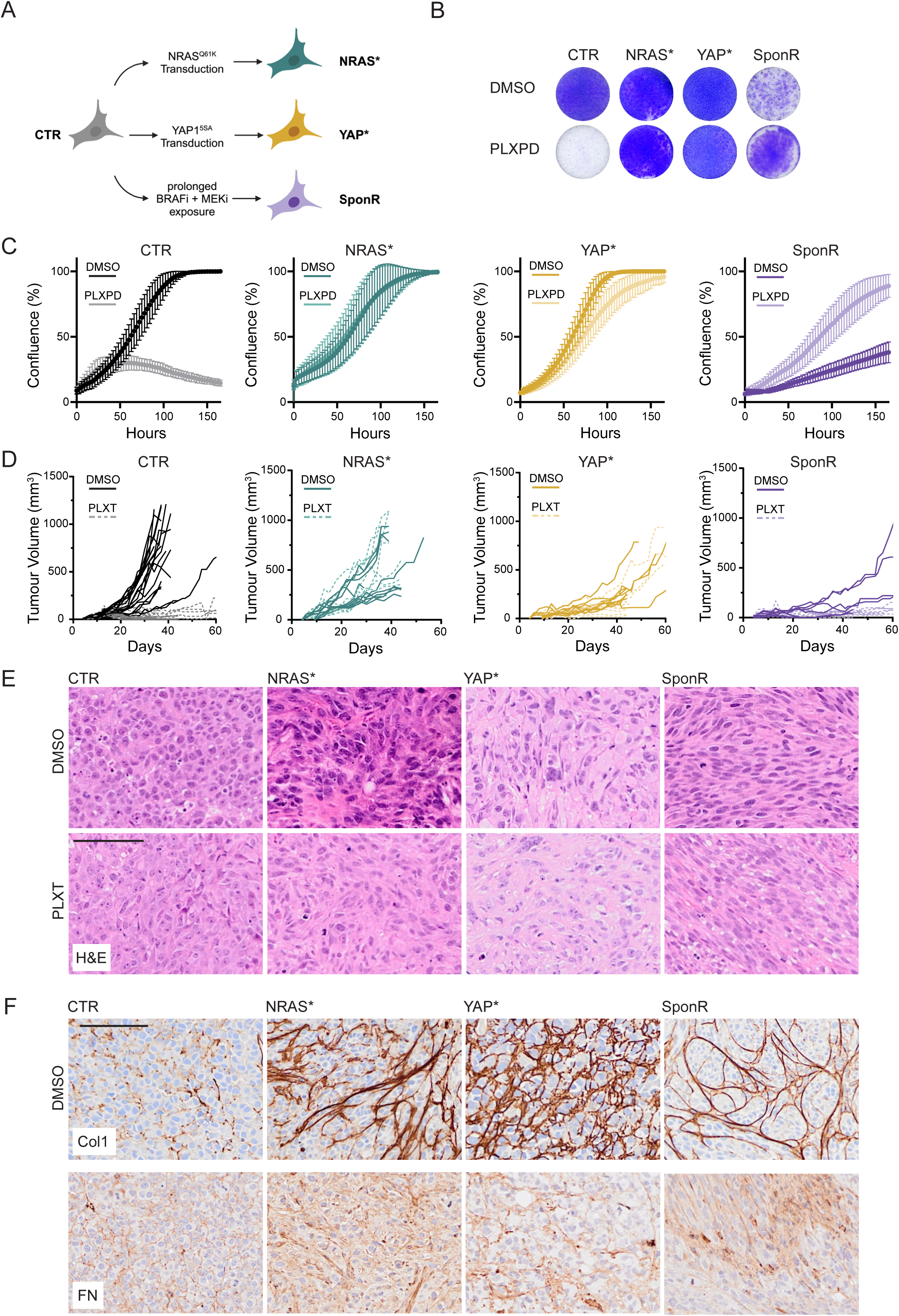
Establishment of three different BRAF/MEKi resistant phenotypes in a single model. A. Schematic illustration of the three different mechanisms by which A375 cells were made resistant to BRAFi and MEKi. B. Crystal violet staining is shown to indicate the growth of the indicated cells cultured with or without BRAFi and MEKi – specifically 1µM PLX4720 and 1µM PD184352 (PLXPD) – 8 days after seeding of cells. C. Plots show the growth of the indicated cells over 160 hours. Lighter shades indicate treatment with 1µM PLX4720 and 1µM PD184352. Data show the mean and standard deviation of three independent replicates. D. Plots show the tumour growth by the indicated cells. Lighter dashed lines indicate mice treated with PLX4720 and Trametinib from day 13 onwards (labelled PLXT and marked with arrow). Each line represents a different tumour/mouse. E. Images show H&E staining of the indicated tumours. Scale bar is 100µm. F. Images show collagen I staining (brown) of the indicated tumours. Scale bar is 100µm.

### Establishment of three different BRAF+MEKi resistant phenotypes in a single model

Distinguishing between these possibilities requires a system in which multiple resistance mechanisms can be generated, compared, and combined within a single isogenic genetic background. We set out to determine if different resistance mechanisms to combined BRAF and MEK inhibition (BRAF+MEKi) push cells into a common resistant state, or if there are multiple distinct therapy resistant states. As both gain of NRAS mutation and transcriptional activation of YAP1 can drive therapy resistance, we engineered A375P (hereafter A375) melanoma cells to contain either mutant NRAS^Q61K^ or constitutively active YAP1^5SA^ using lentiviral transduction (hereafter referred to as NRAS* and YAP* cells, with control cells labelled CTR – Fig. 1A). We additionally cultured A375 cells in BRAF and MEK inhibitors (PLX4720 and PD184352 – hereafter referred to as PLXPD) for 19 weeks until cells had been selected that grew in the presence of the drugs showing spontaneous resistance (hereafter referred to as SponR – Fig. 1A). The ectopic expression of YAP1^5SA^ and NRAS^Q61K^ was confirmed by western blot analysis (Supp. Fig. 2A), with resistance of BRAF+MEKi confirmed by crystal violet assays (Fig. 1B). Intriguingly, the three resistance mechanisms showed markedly different growth dynamics during establishment. A375 cells transduced with NRAS^Q61K^ initially formed much smaller and fewer colonies (Supp. Fig. 2B). Inhibition of BRAF and MEK enhanced the growth of cells transduced with NRAS^Q61K^, suggesting that over-activation of ERK signalling may be responsible for slow growth following the introduction of the active NRAS mutant (Supp. Fig. 2C). Cells transduced with NRAS^Q61K^ grew slowly for at least two weeks, but by 7 weeks their growth was only marginally slower than CTR cells (Supp. Fig. 2D&E). These data are consistent with previous reports of excessive ERK activity leading to growth arrest and the mutual exclusivity of activating NRAS and BRAF mutations in drug-naïve melanoma. In contrast, the introduction of YAP1^5SA^ had minimal effect on the growth rate of A375 cells *in vitro*. In line with previous studies, it took several weeks to select A375 SponR cells that were able to grow in the presence of inhibitors (SponR cells). A375 SponR cells grew slower than YAP* and NRAS* cells in the presence of BRAF+MEKi (Fig. 1C and Supp. Fig. 2F). The growth of SponR cells was even slower in the absence of BRAF + MEKi, indicating that they had a ‘drug addicted’ phenotype as previously shown; this was not observed for NRAS* and YAP* cells (Fig. 1C). Prolonged culture of SponR cells in the absence of BRAF+MEKi led to a phenotypic reversion, with the cells becoming sensitive to BRAF+MEKi once again (Supp. Fig. 2F). This indicates that the SponR cells have not acquired a genetic mutation conferring resistance. Whole exome sequencing confirmed that no known additional mutations linked to BRAF+MEKi resistance were present in the SponR cells, nor the YAP* and NRAS* cells (Supp. Data. 1). As expected, the YAP15SA and NRASQ61K alleles were observed in the YAP* and NRAS* cells, respectively.

**Figure 2.**
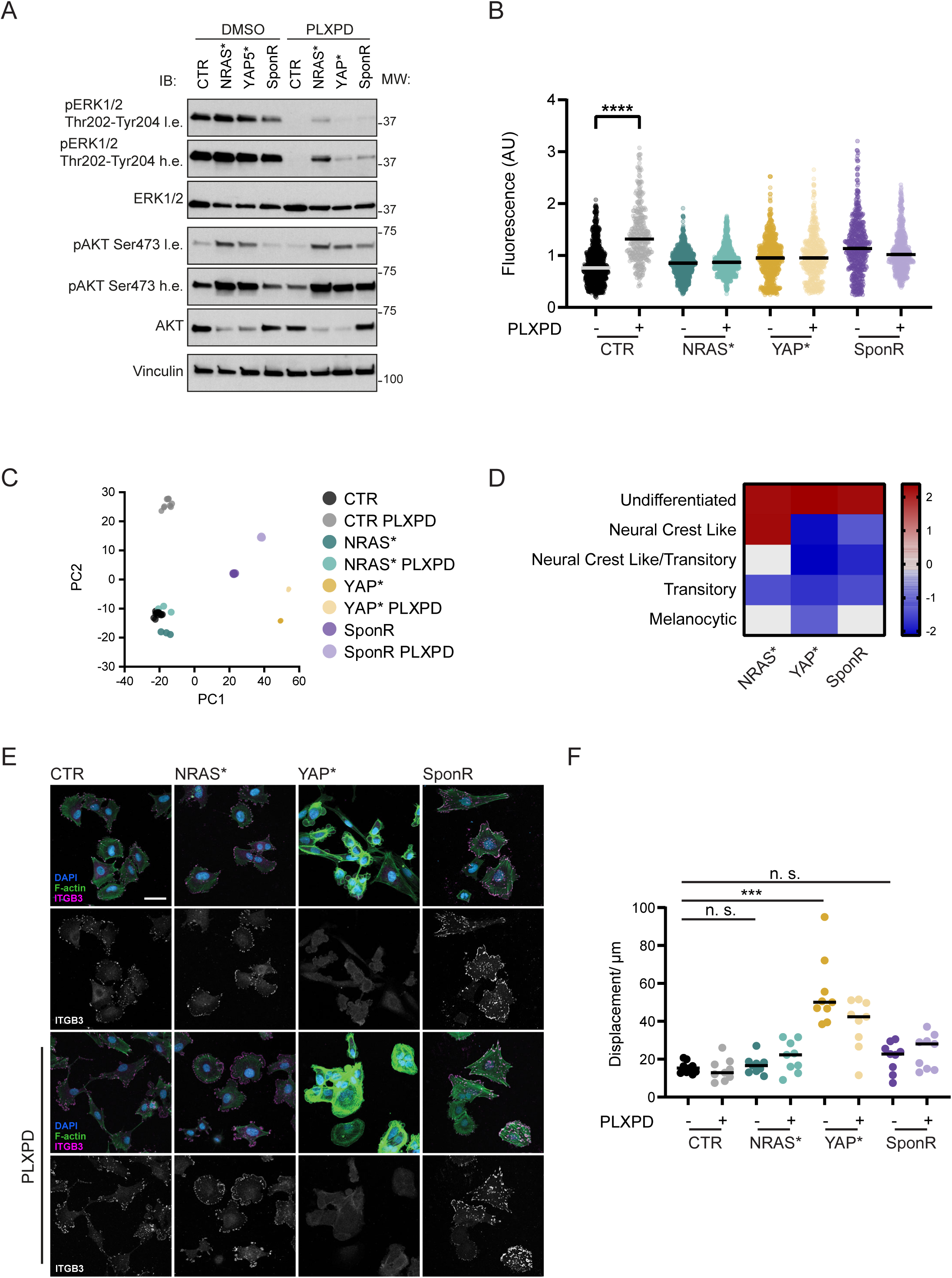
Characterization of three different BRAF/MEKi resistant phenotypes. A. Western blots showing the levels of pERK, total ERK1/2, pAKT, total AKT, and vimentin. Light exposure and heavy exposure are indicated by l.e. and h.e., respectively. Molecular weight markers are shown on the right-hand side. B. Plot shows normalised CellROX staining intensity for the indicated cells with or without 1µMPLX & 1µMPD. Each dot represents a cell with data pooled from three biological replicates. C. Principle component analysis of RNA sequencing of CTR, NRAS*, YAP* and SponR cells cultured with or without 1µMPLX & 1µMPD – labelled PLXPD. Each dot represents a biological replicate, with the size of the dots representing the value of principle component 3, with larger dots indicating a higher value. D. Heatmap shows relative expression of genesets corresponding to the previously described melanoma cell states (Tsoi et al). Red indicates increased expression, while blue indicates reduced expression. E. Images show DAPI (cyan), F-actin (green), and ITGB3 (magenta/white) staining of the indicated cells cultured with or without 1µMPLX & 1µMPD. Images are representative of one of three biological replicates. F. Plot shows the displacement distance in microns over a 24-hour period of the indicated cells cultured with or without 1µMPLX & 1µMPD. Each dot represents an individual field of view with data pooled from three biological replicates.

We next sought to test if A375 NRAS*, YAP*, and SponR cells were resistant to BRAF + MEK targeted therapy *in vivo*. Fig. 1D shows that while control (CTR) A375 tumours were effectively treated with BRAF+MEKi (PLX4720 and Trametinib – abbreviated to PLXT), NRAS* and YAP* tumours were not (see also Supp. Fig. 2G). In addition, the YAP* tumours grew slower (Fig. 1D, Supp. Fig. 2G). The SponR tumours grew slowly, which can be attributed to their addiction to BRAF+MEKi, and the absence of drug treatment in the two-week period following injection. Haematoxylin and Eosin staining revealed that both NRAS* and SponR tumours had increased numbers of spindled morphology cells, while YAP* tumours had a more disorganised architecture with pleomorphic cells and frequent atypical mitoses (Fig. 1E). All three resistance mechanisms were associated with increased collagen I (Col1) deposition, which has previously been linked to therapy resistance and YAP activation (Fig. 1F). Together, the distinct growth kinetics and histological features of NRAS*, YAP*, and SponR tumours confirm that these three resistance mechanisms drive cells into fundamentally different drug-resistant states.

### Phenotypic characterisation reveals divergent signalling, transcriptional, and migratory states

We characterised the phenotypic differences between the YAP*, NRAS*, and SponR cells in more depth. Western blotting confirmed that ERK activity was lower in all resistant cells treated with BRAF+MEKi (Fig. 2A); however, the reduction in NRAS* cells was less pronounced. Activation of PI3K/AKT signalling was observed in NRAS*, YAP* and SponR* cells treated with BRAF+MEKi (Fig. 2A). In line with previous findings (Cesi *et al*., 2017), reactive oxygen species (ROS) were elevated in drug-naïve cells treated with BRAF+MEKi and, to a lesser degree, in spontaneously selected resistant A375 cells (Fig. 2B). However, they were not observed in YAP* cells or NRAS* cells, even when treated with BRAF+MEKi, indicating that these cells have gained mechanisms to avoid oxidative stress upon drug exposure.

To understand better differences among YAP*, NRAS*, and SponR cells, we performed RNA sequencing (RNAseq) in both absence and presence of BRAF+MEKi. NRAS* cells were sequenced shortly after transduction of NRAS^Q61K^ at day 6 (d6) and day 12 (d12), when cells have greatly retarded growth, and after 7 weeks, when cells were growing at equivalent rates to empty vector transduced cells (Supp. Fig. 2D). Both gene ontology and principal component analysis confirmed that the three resistance mechanisms reflect different states (Fig. 2C and Supp. Fig. 3A). Moreover, the three resistance mechanisms differed in their expression of the markers used to demonstrate heterogeneity in cell state in patient samples, indicating that our models reflect clinically relevant diversity in cell state (Supp. Fig. 3B). Control A375 cells are most similar to a neural crest-like state (Wouters *et al*., 2020). NRAS* cells (growing at normal rate 7 weeks post-transduction) were even closer to this neural crest-like state and also showed some patterns of gene expression linked to development of the nervous system and undifferentiated cell state (Supp. Fig. 3A). Interestingly, twelve days post-transduction with NRAS^Q61K^, NRAS* cells expressed high levels of the inflammatory cytokines IL1A, IL1B, and TNF that then returned to baseline levels seven weeks post-transduction (Supp. Fig. 3C). The relevance of this observation was confirmed in functional experiments, which demonstrated that TNFα and IL1β selectively inhibit the growth of NRAS* cells (Supp. Fig. 3D). This transient inflammatory response may act as an initial brake on NRAS* cell expansion, consistent with the slow colony formation observed during establishment (Supp. Fig. 2B) and suggests that NRAS-mutant cells must resolve this autocrine inflammatory programme before achieving stable resistance. YAP* and SponR cells were in undifferentiated/mesenchymal-like states but did not show increased neural crest gene expression (Fig. 2D). They also had similar changes in the expression of genes previously linked to BRAF inhibitor resistance (Supp. Fig. 3E). Despite these commonalities, YAP* and SponR cells were in distinct states (Fig. 2C). SponR cells showed altered expression of negative regulators of signal transduction (Supp. Fig. 3A), which is concordant with previous findings (Wang *et al*., 2018; Das Thakur *et al*., 2013), while YAP* cells had particularly prominent changes in expression of genes related to cell migration (Supp. Fig. 3A).

**Figure 3.**
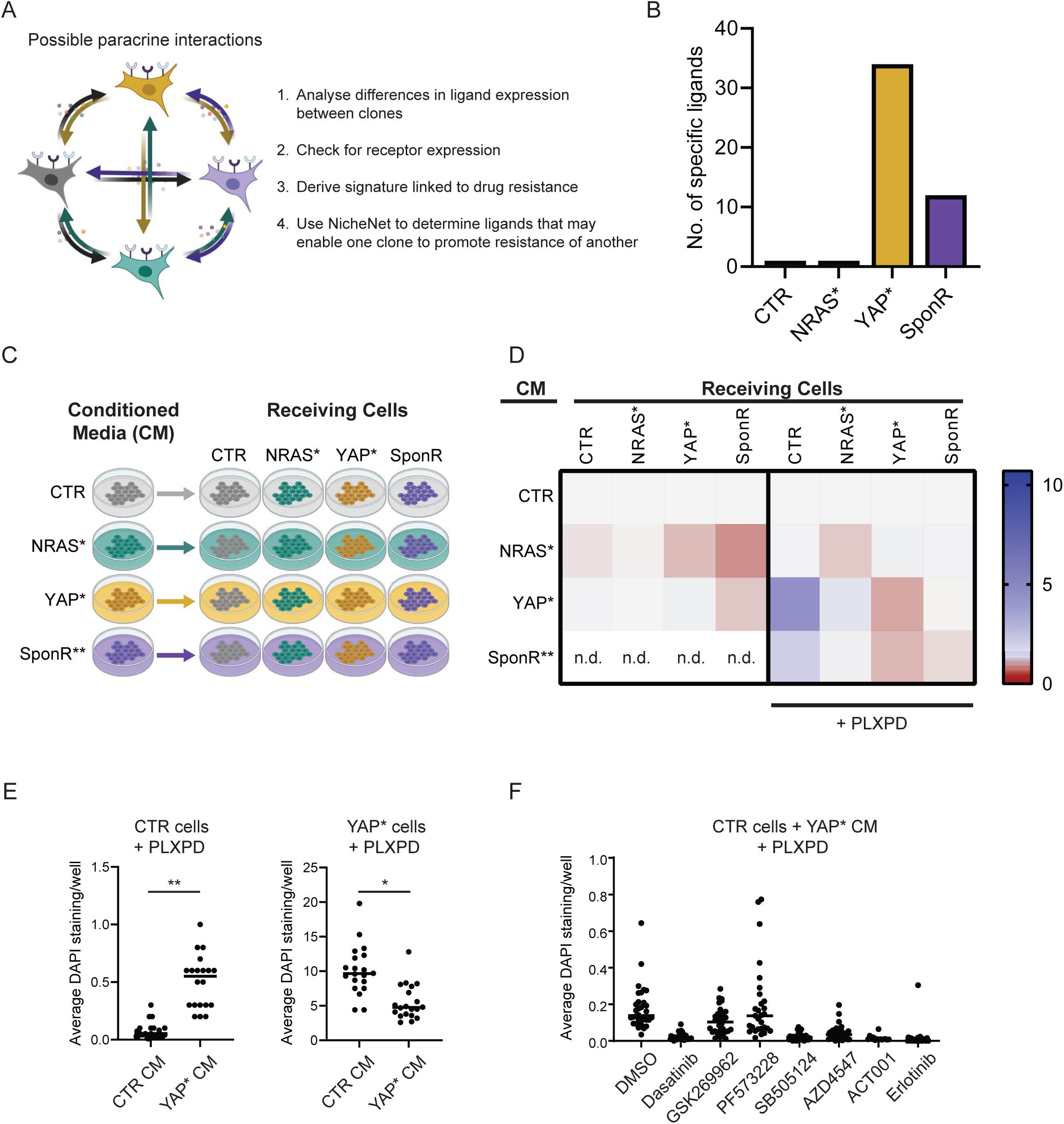
Co-cultures reveal inter-subclone crosstalk. A. Scheme underpinning NicheNet analysis is shown. B. Plot shows the number of unique ligand signals as inferred using NicheNet analysis generated by CTR, NRAS*, YAP*, and SponR cells C. Schematic - Schematic diagram illustrates the collection of conditioned media from different A375 lines and transfer onto different A375 lines in the absence or presence of 1µM PLX4720 + 1µM PD184532. Note SponR conditioned media contains residual PLX and PD from the routine culture of these cells in the presence of drug. D. Conditioned media experiments - Heatmap shows the fold effect of different conditioned media relative to media conditioned by parental A375 cells. Blue indicates boosted cell numbers and red indicates reduced cell numbers in the presence of the indicated conditioned media (NRAS*, YAP*, or SponR). n.d. denotes not determined due to the confounding presence of BRAF and MEK inhibitors in the conditioned media from SponR cells. Scale reflects fold change relative to control conditioned media. E. Plots show the relative number of cells, as determined by DAPI staining, in either CTR or YAP* cells cultured in the presence of 1µM PLX & 1µM PD and either CTR or YAP* conditioned media (CM). Each dot represents a field of view with data pooled from three biological replicates. F. Plots show the relative number of cells, as determined by DAPI staining, for CTR cells cultured in YAP* conditioned media (CM) in the presence of 1µM PLX4720 & 1µM PD184352 and the inhibitors indicated. Each dot represents a field of view with data pooled from two biological replicates.

The analyses above prompted us to explore if NRAS*, YAP*, and SponR cells had different morphologies or migratory properties. YAP* cells, but not NRAS* or SponR cells, had an elongated morphology and fewer ITGB3 positive focal adhesions – both in DMSO and combined BRAF+MEKi conditions (Fig. 2E). Transcriptional analysis indicated that ITGB3 and many other integrins were down-regulated in YAP* cells (Supp. Fig. 3F). YAP* cells also had features associated with changes in migratory behaviour, including plasma membrane blebs and atypical nuclear morphology. Fig. 2F shows that the displacement of YAP* cells over 24hrs (distance between position at start and end of the 24hr period) was almost four times higher than that of CTR, NRAS*, and SponR cells.

Together these data show that the same starting population of melanoma cells can be driven into three highly distinctive drug-resistant states, exhibiting clear differences in transcriptional programmes, handling of ROS, addiction to BRAF+MEKi, cell and nuclear morphology, and cell migration. This diversity within a single genetic background implies that any therapeutic strategy must account for multiple co-existing resistant states, not just the dominant one.

### Crosstalk between resistant states reveals that YAP1-high cells provide paracrine support while inhibiting their own expansion

Analysis of the transcriptional data indicated that several molecules involved in cell-cell signalling were altered in the resistant states, such as AXL and EGFR (Supp. Fig. 3E), raising the possibility of functional communication between co-existing resistant states. To explore this possibility, we performed NicheNet analysis (Fig. 3A). We examined if cells in the different resistant states increased the production of ligands that might drive transcription programmes linked to continued growth in the presence of BRAF+MEKi. We additionally required that the receptor must be expressed by the recipient cell. This revealed that YAP* cells expressed the greatest number of ligands capable of crosstalk to other cell states (Fig. 3B), including modulators of TGFβ signalling – TGFB2, BMP4, FST, tyrosine kinase ligands – PDGFB, NRG, GAS6, various ECM components – COL1A1, COL5A1, COL12A1, COL13A1, and modulators of Wnt signalling – WNT5B, WNT7A, DKK1, (Supp. Fig.S4A&B). These analyses prompted us to explore possible functional interactions between the different resistant states. CTR, YAP*, NRAS*, and SponR cells were cultured in conditioned media from each different state, both in the presence and absence of BRAF+MEKi (Fig. 3C shows experimental scheme). The results show that conditioned media from YAP* cells supports the growth of CTR and NRAS* cells in the presence of BRAF+MEKi (Fig. 3D & E). Intriguingly, YAP* conditioned media inhibits the growth of YAP* cells (Fig. 3D & E), demonstrating that YAP-high cells simultaneously support other resistant states while restraining their own expansion. This self-inhibitory effect is consistent with the slower growth rate of YAP* tumours *in vivo* (Fig. 1D and (Zhang *et al*., 2020)) and provides a mechanism by which heterogeneous tumours could maintain a stable mixture of cell states rather than converging on a single dominant clone. In contrast, conditioned media from NRAS* and SponR* cells did not significantly alter the growth of other cell states (Fig. 3D & E), indicating that the supportive paracrine function is specific to YAP*.

Having determined that YAP* cells produce supportive paracrine signals, we sought to block them using inhibitors targeting pathways prominently represented in the NicheNet analysis. CTR cells were cultured in the presence of supportive signals from YAP* cells, but with the addition of inhibitors blocking receptor tyrosine kinases and downstream signalling (EGFR, AXL, PDGFR, PI3K, Src-family kinases), TGFβ signalling, and integrin/ECM signalling (Cilengitide, FAK and ROCK/Rho-kinase). Figure 3F shows that Src-family kinases, TGFβR, PAI-1, and EGFR are required for the receipt of supportive signals from YAP* cells, while FAK and ROCK kinases were not. These data demonstrate that YAP* cells engage in paracrine signalling via multiple mediators that limit the efficacy of BRAF+MEKi on sensitive drug naïve cells.

### Agent-based modelling predicts that enhanced migration favours YAP5SA cells, but crosstalk prevents their total dominance

The experimental data above reveal two competing forces: YAP* cells have a migration advantage that favours their expansion, but their paracrine support of other cell states and self-inhibition could prevent competitive exclusion. To predict how these forces interact over time and in the context of therapy, we implemented an agent-based model of tumour growth (Fig. 4A&B). In brief, every cell is represented by an agent with the potential to proliferate, die, or move. In addition, cells can signal to other cells in the tumour, with YAP* cells providing signals that protect CTR cells from drug, which mirrors our experimental observation (Fig. 3A-E). The model is initiated with a single cell that then grows into a tumour colony. The properties of the initiating cell reflect the drug naïve CTR A375 cells, in terms of migration, proliferation, and response to the addition of BRAFi+MEKi (Supp. Table 2). The relatively slow migration of A375 cells means that tumour colonies develop high cell density at the centre, which reduces proliferation in this zone. Supp. Fig. 5A provides experimental confirmation of this phenomenon. During the simulation, cells have a probability of transitioning to other states reflecting YAP*, and NRAS*. We did not model the transition to the SponR state because it happens with a very low frequency, both *in vitro* and *in vivo*. YAP* cells have increased migratory behaviour (based on Fig. 2F) and a slower proliferation rate (based on Figs 1D and 3D). NRAS* and SponR cells initially have a slow growth rate, which reflects the slow proliferation rate observed after the introduction of NRAS^Q61K^ and initial exposure to BRAFi+MEKi, respectively (based on Fig. 2 and Supp. Fig. 3). Figure 4C shows examples of the model output when the rate of transition to NRAS* and YAP* states is low (see also Supp. Movies 1&2). In the absence of simulated drug, tumours grow rapidly with YAP* cells only contributing a small amount to the overall tumour (indicated by point (1) in Fig. 4C). YAP* cells are slightly favoured compared to CTR cells due to their enhanced migratory behaviour (Supp. Fig. 5C), although if the difference is too great then the advantage is lost. When the addition of drug is simulated, tumours initially shrink, entering a drug tolerant state, before re-growing as drug resistant tumours (indicated by points (2) and (3) in Fig. 4C). These transitions are associated with an increase in the proportion of YAP* cells. The supportive effect of YAP* cells on CTR cells means that they do not dominate totally, with a greater boosting effect leading to a greater proportion of CTR cells (Supp. Fig. 5D). The enhanced migratory capability of YAP* cells is predicted to further enhance their advantage (Supp. Fig. 5D). Simulations with a high transition rate to YAP* and NRAS* states (p=0.0001) exhibited only limited responses to therapy. If the transition rate between states is low (e.g p=0.00003), then therapy initially reduces the tumour size, with YAP* cells predicted to be more prominent in the residual tumour (Supp. Fig. 5E). NRAS* cells only make a minor contribution due to their initial low growth rate (Supp. Fig. 2B).

**Figure 4.**
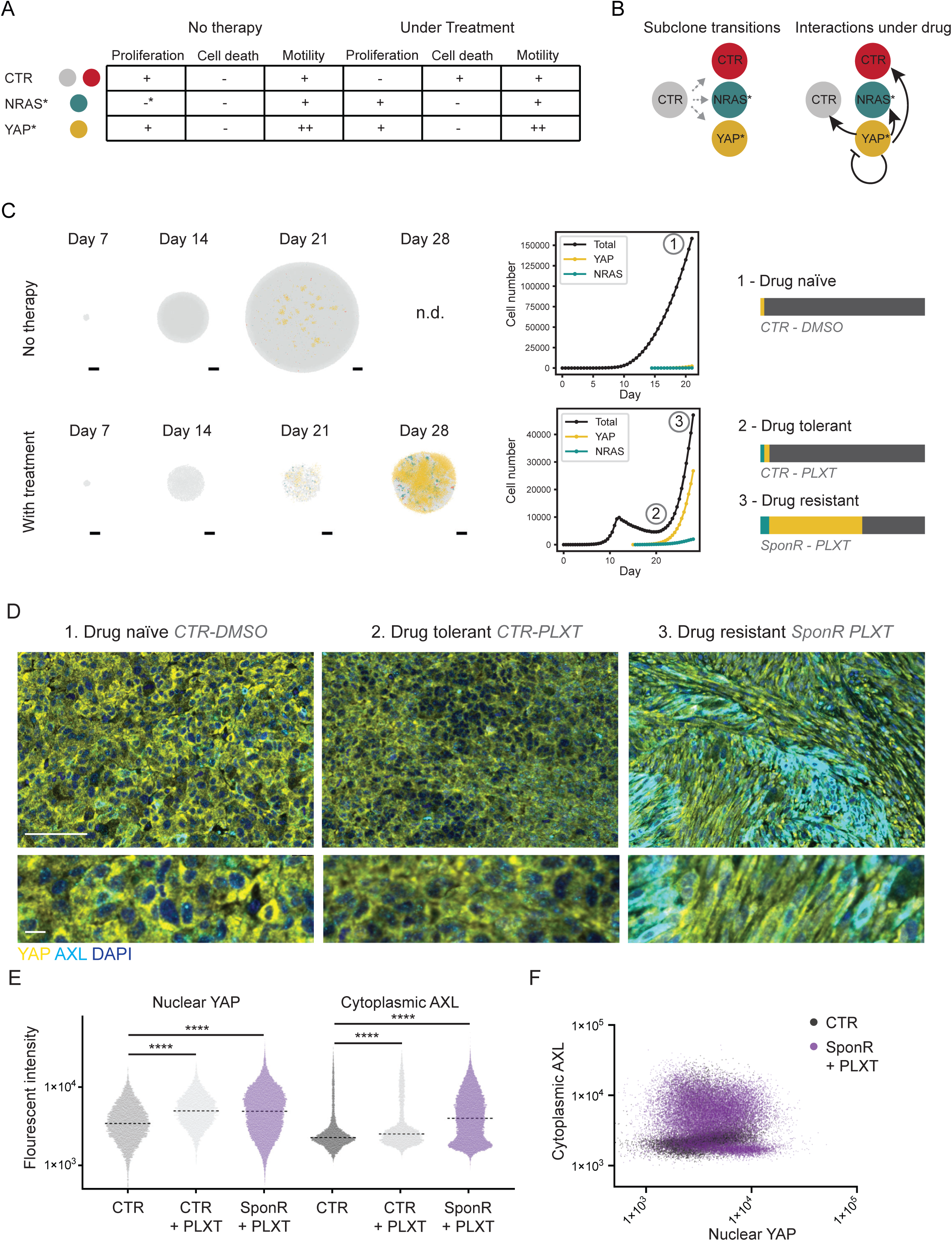
Computational modelling predicts tumour heterogeneity as favoured state following therapy. A. Table summarising the features of CTR, NRAS*, and YAP* cells is shown. B. Left: schematic illustration of transitions permitted between CTR (grey – no transition, red – no transition but labelled for comparison with other transitions), NRAS* (dark green), and YAP* (yellow) cells. Right: schematic illustration of the autocrine and paracrine effects between CTR (grey – no transition, red – no transition but labelled for comparison with other transitions), NRAS* (dark green), and YAP* (yellow) cells. Transitions are indicated with grey dashed arrows and autocrine and paracrine interactions with black arrows. C. Representative images of model outputs reflecting no treatment (upper panels) and with treatment started on day 11 (lower panels). CTR cells are either grey – no transition, or red – no change in cell state but labelled for comparison with other transitions, NRAS* cells are dark green, and YAP* cells are yellow. Right-hand plots show the total cell number (black), NRAS* cell number (dark green), and YAP* cell number (yellow). Key points are indicated with (1) maximal growth of drug-naïve tumour, (2) Residual disease in drug tolerant tumour, and (3) maximal re-growth in drug resistant tumour. Bar plots summarise the cellular composition at points (1), (2), and (3). D. Images show staining of CTR, CTR + BRAFi+MEKi, and SponR + BRAF+MEKi tumours reflecting (1) drug naïve, (2) drug tolerant, and (3) drug resistant tumour states, respectively. DAPI, YAP1, and AXL are shown in blue, yellow, and cyan, respectively. Scale bars are 100µm (upper panels) and 10µm (lower panels). E. Quantification of nuclear YAP levels (left) and cytoplasmic AXL levels (right) in control drug-naïve (CTR), drug-tolerant disease (CTR + PLXT), and drug resistant disease (SponR + PLXT). >8000 cells from >5 tumours quantified for each condition. F. Plot shows nuclear YAP and cytoplasmic AXL levels in control drug-naïve (CTR, dark grey) and drug resistant disease (SponR + PLXT, magenta). >8000 cells from >5 tumours quantified for each condition.

**Figure 5.**
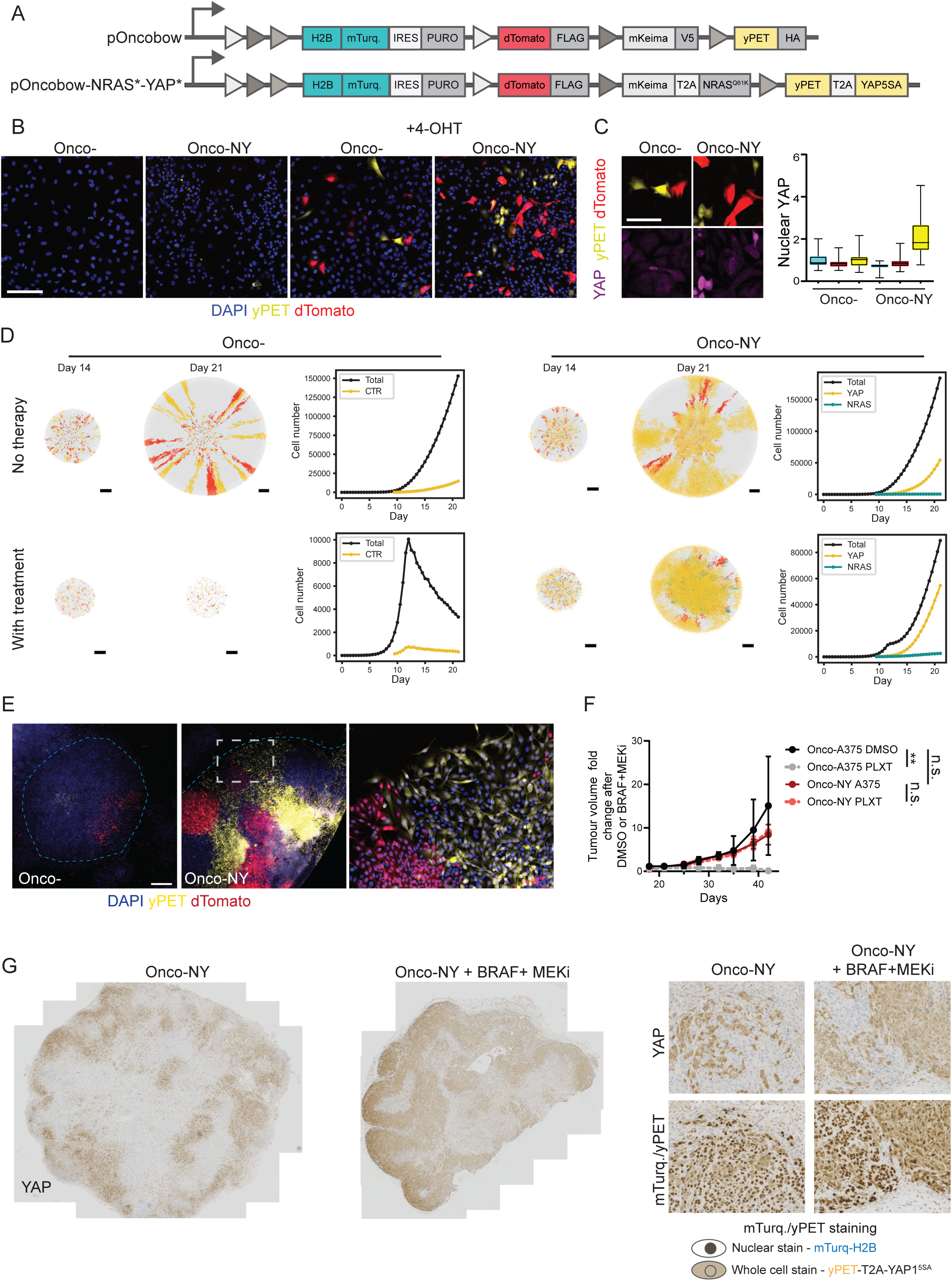
An experimental model of intra-tumour heterogeneity. A. Images show schematic representations of Oncobow cassettes. Different coloured triangles represent different loxP sequences. B. Images show DAPI staining (blue), tdTomato expression (red), and yPET expression (yellow) in A375 cells with stably integrated Cre-ERT and Oncobow or Oncobow-NY cassettes cultured in either the absence or presence of 20nM 4-OHT for 72 hours. Scale bar is 200 microns. C. Images show YAP1 staining (magenta), tdTomato expression (red), and yPET expression (yellow) in A375 cells with stably integrated Cre-ERT and Oncobow or Oncobow-NY cassettes cultured in either the absence or presence of 20nM 4-OHT for 72 hours. Scale bar is 100 microns. D. Panels shown model output with following synchronised timing of transitions to NRAS* and YAP* states, mirroring the Onco- and Onco-NY experimental setting. Left side shows no drug treatment, right side shows with simulated drug treatment. CTR cells are either grey – no transition or red – no change in cell state but labelled for comparison with other transitions, NRAS* cells are dark green, and YAP* cells are yellow. Graphs show the total cell number (black), NRAS* cell number (dark green), and YAP* cell number (yellow). E. Images show an Onco-A375 colony (left) and an Onco-NY-A375 colony (mid), with a zoomed image of the edge of the Onco-NY-A375 colony shown on the right. DAPI is shown in blue, yPET in yellow, and tdTomato in red. Dashed fine blue line indicates colony edges. Scale bar is 500µm. F. Plot shows the fold change in tumour volume following administration of PLX4720 + Trametinib or DMSO control in Onco-A375 or Onco-NY-A375 cells. Four mice per group with mean and standard deviation shown. G. Images show location of high YAP1 expressing cells in representative Onco-NY tumours either DMSO treated (left) or PLX4720 and Trametinib treated (right). Higher magnification images are shown on the right along with the corresponding region of a serial section stained for CFP/yPET. Nuclear signal indicates H2B-mTurq expression in non-recombined cells and whole cell signal indicates yPET signal in cells expressing YAP1^5SA^.

The analyses described above predict that YAP* cells will not completely dominate and that heterogeneity in YAP activity will be the optimal state for melanoma. To explore this prediction experimentally, we stained the tumours described in Figure 1 for YAP1 and AXL – a marker of mesenchymal cell state (Fig. 4D). CTR tumours reflect the growing drug naïve state indicated by point (1) in Fig. 4C. In line with the model’s prediction, these tumours had low levels of nuclear YAP1, with the majority of the protein localised in the cytoplasm, and low levels of AXL (Fig. 4D&E). The residual disease following treatment of CTR tumours with BRAF+MEKi, which reflects the drug tolerant state in Fig. 4C, showed more variable YAP levels. Most strikingly, SponR tumours generated with cells that had undergone a prolonged selection for growth in the presence of BRAF+MEKi showed pronounced heterogeneity in nuclear YAP levels and AXL expression (Fig. 4D right panels and Fig. 4E). Combined analysis of nuclear YAP1 and AXL clearly indicated the emergence of two cell states in the tumours capable of growing in BRAF+MEKi (Fig. 4F). Interestingly, nuclear YAP1 and AXL levels were not correlated, suggesting the presence of other mechanisms besides YAP1 in driving AXL expression. Overall, these analyses are concordant with the *in silico* predictions and indicate that therapy selects for tumours containing heterogeneous cell states, and not a single dominant state.

### An experimental model of subclonal heterogeneity

The preceding experiments established that resistant subclones engage in cooperative signalling and that computational modelling predicts stable co-existence rather than competitive exclusion. However, because each resistant state was generated and cultured independently, these data cannot resolve whether stable heterogeneity emerges spontaneously when resistance mechanisms arise within the same tumour. In patients, the mutational processes and cell state transitions that generate sub-clonal heterogeneity occur in the context of an established tumour, not in isolation. We, therefore, sought to build a system that recapitulates this by allowing resistant states to emerge within a pre-existing tumour, while maintaining experimental control over the rate of transition between states. To this end, we adapted a ‘Brainbow’ cassette (hereafter called Oncobow – Bhargava et al., bioRxiv) to include NRAS^Q61K^ and YAP1^5SA^ transgenes and introduced this into A375 cells expressing CreERT2. Upon Cre-mediated recombination, individual cells stochastically express one of three cassettes: YAP1^5SA^ coupled to the fluorophore yPET, NRAS^Q61K^ coupled to Keima, or tdTomato as a neutral control. Figures 5A & 5B show the construct design and confirmation of tamoxifen inducible recombination, with only a low level of recombination in the absence of tamoxifen. Following recombination of Oncobow-NRAS*- YAP* (hereafter referred to as Onco-NY A375), but not the empty Oncobow cassette (hereafter referred to as Onco-), yPET expressing cells had elevated levels of YAP1 (Fig. 5C). Genomic PCR indicated that the NRAS locus recombined as expected (Supp. Fig 6A).

**Figure 6.**
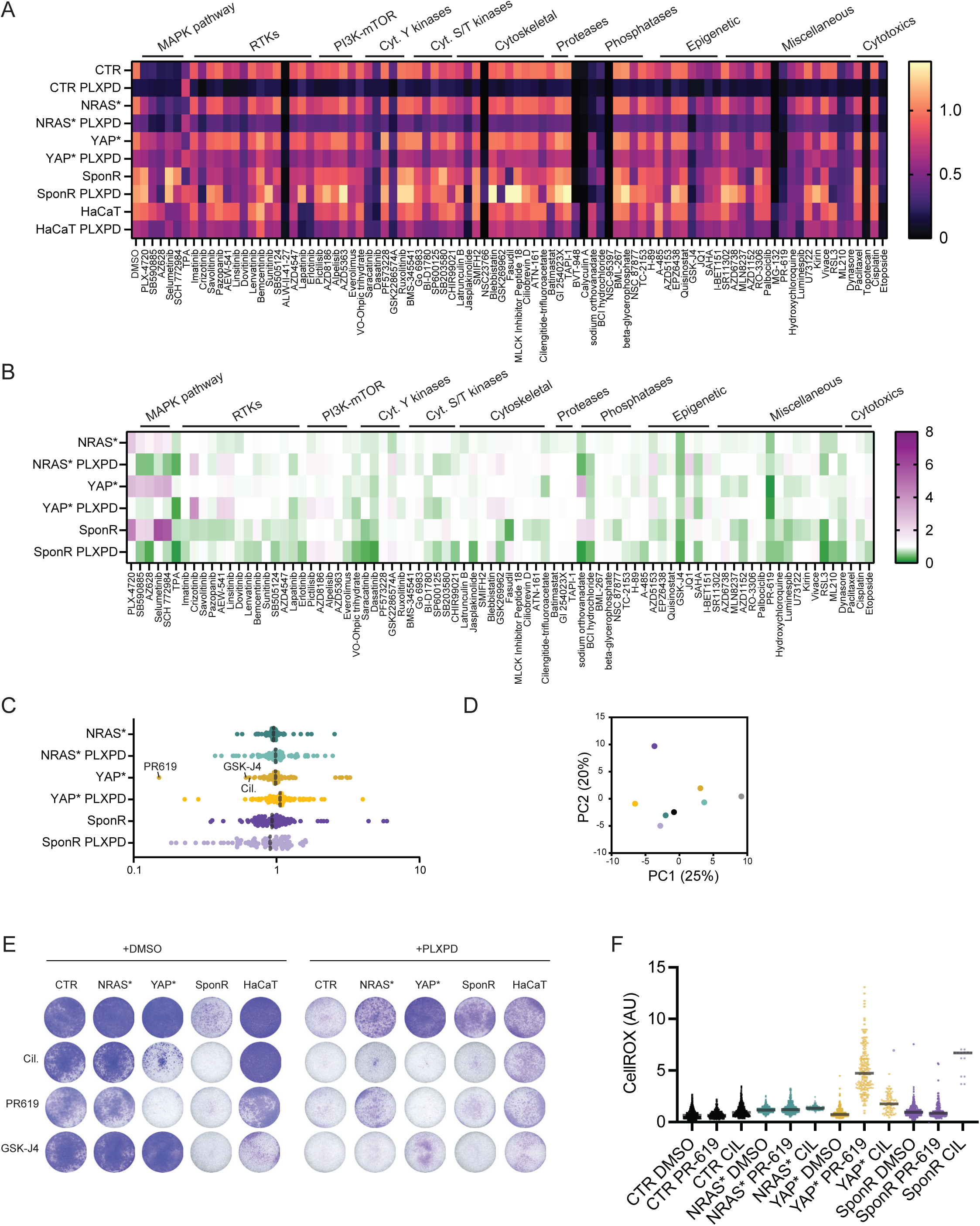
Identification vulnerabilities of different drug resistant states. A. Plot shows the effect on cell viability measurements (CellTiter-Glo) of the indicated compounds on CTR, NRAS*, YAP*, and SponR cells in the presence or absence of 1µM PLX4720 and 1µM PD184532 after 72 hours. Red indicates lower viability values relative to the cells in the absence of any drug, blue indicates higher viability values. Data shown are the mean of three biological replicates. B. Plot shows whether the sensitivity of NRAS*, YAP*, and SponR cells to the indicated compounds in either the presence or absence of PLX PD is increased (green shading) or decreased (magenta shading) relative to CTR cells. Cell viability was determined by CellTiter-Glo assay after 72 hours of drug exposure. Data shown are the mean of three biological replicates. C. Plots shows the spread of values generated in the differential sensitivity analysis shown in (B) with strongest vulnerabilities of YAP* cells highlighted. D. Principal Component Analysis plot indicating the relative similarity of the drug sensitivity profiles of CTR, NRAS*, YAP*, and SponR cells. Numbers in brackets are the percent of variance explained by the principal component. E. Images show crystal violet staining of the indicated cells treated with or without 1µM PLX4720 & 1µM PD184352 and the indicated compounds at the following concentrations. One of three representative experiments is shown. F. Plot shows normalised CellROX staining intensity for the indicated cells with or without 5µM PR619 or 10µM Cilengitide. Each dot represents a cell with data pooled from three biological replicates.

We adopted an integrated *in silico* and experimental strategy to explore the behaviour of our model with inducible intra-tumour heterogeneity. Simulations were initiated with the rate of transition between cell states set to zero, the transition was then increased to mimic tamoxifen-induced recombination at a time point corresponding to 7 days (Fig. 5D). This regimen led to the prediction that cells expressing YAP5SA would be disproportionately represented compared to those expressing tdTomato, particular at the frontier of tumours (right hand panel - Fig. 5D and Supp. Movies 3-6). This prediction is based on the increased migration of these cells enabling them to move from less proliferative zones in the tumour centre to the tumour edge. To test this, we seeded A375 cells as single cells before triggering recombination at 120 hours and allowing growth until millimetre scale colonies containing >10^4^ cells had formed. Notably, YAP* expressing sub-clones were frequently observed expanding at the colony edge and were favoured compared to dTomato sub- clones in the same colony and compared to yPET sub-clones in Oncobow cells (Fig. 5E), which confirms the model prediction that increased migration confers a selective advantage in contexts when space for growth is limited.

We then proceeded to explore the behaviour of our heterogeneous tumour model *in vivo*. Onco-A375 and Onco-NY A375 cells were injected into the flanks of mice, with tamoxifen administered to drive recombination and diversity of cell states prior to treatment with PLX and Trametinib. Figure 5F shows that while Onco-A375 tumours responded well to BRAF+MEK inhibition, Onco-NY A375 tumours were refractory to therapy. Histological analysis confirmed that high YAP1 expression was abundant around the edges of Onco-NY A375 tumours (Fig. 5G). Onco-NY tumours also exhibited the altered nuclear morphologies observed in the clonal YAP* tumours (Supp. Fig. 6B and Fig. 1E). Staining with a pan- yPET/GFP/CFP antibody was able to discriminate non-recombined H2B-mTurquoise (CFP derivative) cells from yPET/YAP cells on the basis of nuclear vs whole cell staining. Figure 5G highlights the presence of non-recombined cells even in the presence of drug and the concordance of yPET and YAP1 staining. These data confirm that although YAP* cells have a migration advantage, tumours reach a stable equilibrium containing multiple co-existing cell states rather than converging on YAP* dominance. Strikingly, the tumours with sub- clonal YAP1^5SA^ did not exhibit the growth disadvantage clearly evident in tumours with clonal YAP1^5SA^ (c.f. Fig. 1D and Fig. 5F), indicating that tumours composed of multiple co-existing cell states outperform those dominated by a single resistant state.

### Distinct vulnerabilities of different BRAF+MEKi resistant states

To identify strategies for treating tumours with multiple concurrent resistance mechanisms, we mapped the sensitivity of CTR, NRAS*, YAP*, and SponR cells to a targeted library of signal transduction inhibitors and chemotherapies, including those used in Fig. 3. To estimate the therapeutic window, we also screened the effect of the compounds on non-transformed HaCaT skin cells (Fig. 6A). Fig. 6A shows a heatmap of the effect of the compounds on cell viability, as measured by ATP levels. The analyses confirmed NRAS*, YAP*, and SponR cells all had reduced sensitivity to a range of inhibitors targeting RAF, MEK, and ERK signalling (note the lighter shade of red Fig. 6A and Supp. Fig. 7A). Interestingly, AKT inhibition using AZD5356 (Capivasertib) did not affect the viability of the NRAS* and YAP* cells, indicating that the increased AKT activity (Fig. 2A) in these cells was not functionally relevant (Supp. Fig. 7A). AXL inhibition using Bemcentinib also had minimal effect on NRAS* and YAP* cells (Fig. 6A and Supp. Fig. 7A). Other previously reported strategies targeting HDACs, Bromodomain proteins, or GPX4 were not effective across all resistance mechanisms (Supp. Fig. 7A). Compounds that were effective at eliminating the resistant cells, but were similarly or more toxic to HaCaT cells were not pursued further (indicated by similar shading Fig. 6A e.g., dasatinib and MG-132). To identify compounds that might represent new acquired vulnerabilities able to target the resistant cells, we replotted the data comparing the viability of CTR cells, with or without BRAF+MEKi, to the NRAS*, YAP*, and SponR cells (Fig. 6B). SponR cells had the largest number of new vulnerabilities compared to CTR (indicated by green shading in Fig. 6B and the greater number of points less than 1 in Fig. 6C). In contrast, NRAS* and YAP* cells acquired rather few additional vulnerabilities compared to the CTR A375. Moreover, there were almost no common drug vulnerabilities across all three resistance mechanisms. PCA of the drug sensitivity of the CTR and resistant cell lines revealed that each resistance mechanism had distinct changes in vulnerability to the compounds tested, with NRAS* cells having a profile similar to CTR cells and YAP* and SponR cells having more divergent profiles (Fig. 6D).

**Figure 7.**
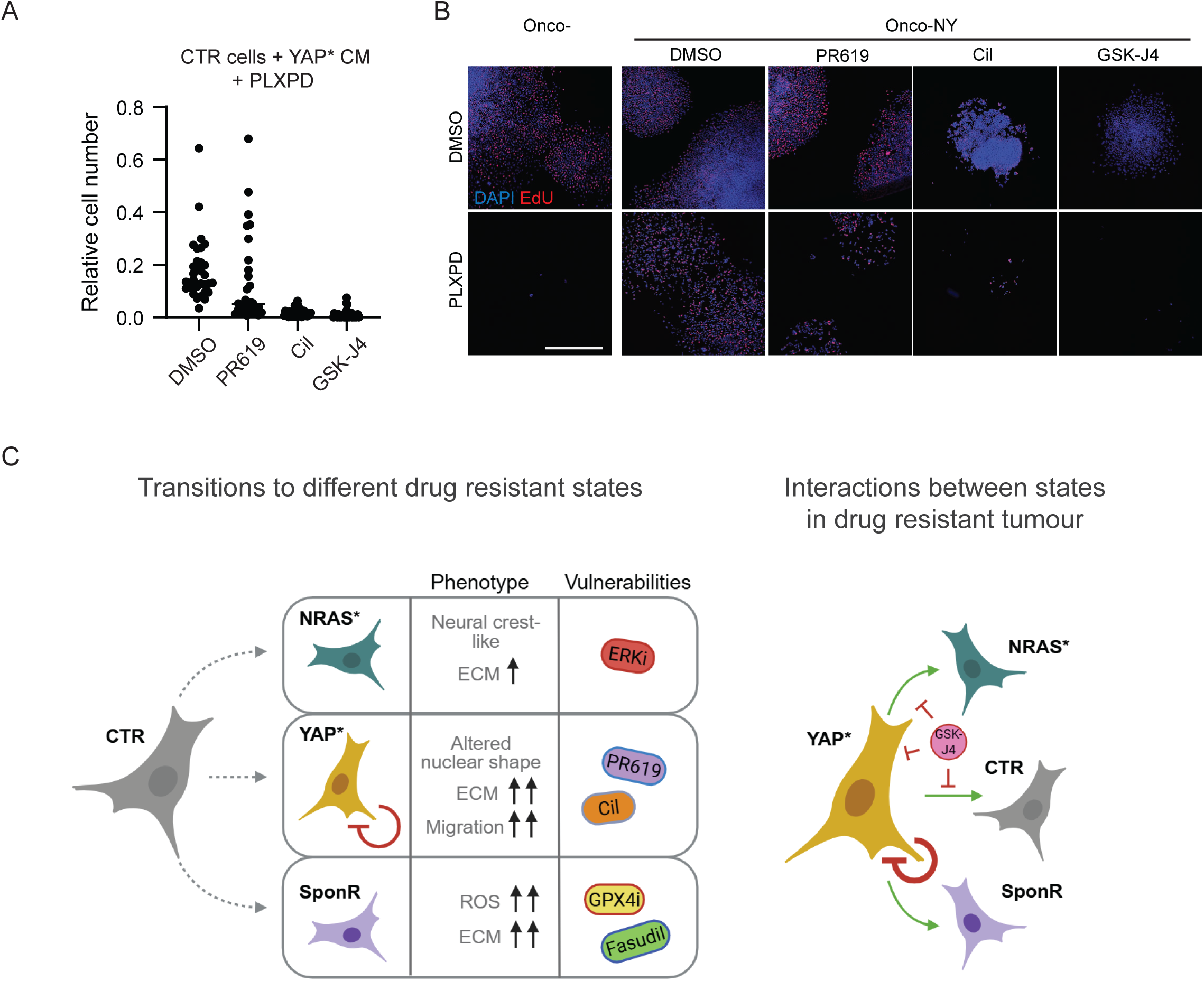
Targeting multiple resistance mechanisms simultaneously. A. Plots show the relative number of cells, as determined by DAPI staining, for CTR cells cultured in YAP* conditioned media (CM) in the presence of 1µM PLX4720 & 1µM PD184352 and the inhibitors indicated. Each dot represents a field of view with data pooled from three biological replicates. B. Images show nuclei stained with DAPI (blue) and EdU (red) of Onco-NY-A375 cells following 4-OHT treatment (all wells) and treatment with the indicated compounds. Scale bar is 1mm. C. Illustration summarises three different BRAF+MEKi resistant states (left) and how they form an ecosystem that favours continued heterogeneity in YAP1 activity (right)

Given the role of the YAP high state in conferring resistance to other cells states (Fig. 3D&E) and its ability to act as a super-competitor (Figs 4&5), we homed in on compounds that were most effective at targeting this cell state. Notably, the DUB inhibitor PR619 (Altun *et al*., 2011), the integrin-engaging peptide Cilengitide and the histone lysine demethylase inhibitor GSK-J4 (Dalpatraj, Naik and Thakur, 2023) were highly effective against YAP* cells. The sensitivity of YAP* cells to GSK-J4 is notable because it inhibits the H3K27 demethylases JMJD3 and UTX, suggesting that active demethylation of H3K27 is required to maintain the YAP*-associated transcriptional state. We next tested the efficacy of these compounds in long term crystal violet assays, which can have slightly different results from MTT assays (Śliwka *et al*., 2016). This revealed PR619 and Cilengitide to be highly effective against YAP* cells as single agents, but GSK-J4 showed a smaller effect than in the MTT assays (Fig. 6E). All compounds were effective in combination with BRAF+MEKi. We attribute the efficacy of Cilengitide to the low levels of ITGB3 and cell matrix adhesions in YAP5SA cells rendering them highly sensitive to any further reduction in the functionality of RGD-binding integrins (Fig. 2E). To try to understand why YAP* cells might be sensitive to these compounds we performed ROS analysis. This revealed that PR619, but not Cilengitide, increased the levels of ROS in YAP* cells (Fig. 6F), leading to the conclusion that the ability of YAP* cells to manage the oxidative stress that would be induced by BRAF+MEKi depends on the activity of de-ubiquitinating enzymes. Similar sensitivities to PR619 and Cilengitide were observed in a second model of YAP-mediated resistance to targeted therapy, specifically resistance to third generation EGFR inhibitors in lung cancer (Supp. Fig. 7B). Collectively, these data reveal that the three resistance mechanisms have largely non-overlapping drug sensitivities, underscoring the challenge of treating heterogeneous resistant tumours with any single agent. However, they also identify a small number of compounds, notably GSK-J4 and Cilengitide, that are selectively effective against the YAP* state, which our earlier data identify as the central node sustaining inter-subclone cooperation.

### Effective targeting of multiple co-existing resistant melanoma states

Our data predict that effective treatment of heterogenous resistant tumours requires simultaneous targeting of the YAP-high state and the inter-subclone signalling that sustains other resistant cells. To test this directly, we compared the effect of targeting either YAP* cells, crosstalk between subclones, or both YAP* cells and crosstalk. We investigated which compounds are able to hinder the supportive signal emanating from YAP* cells to CTR cells. Figure 7A shows that both Cilengitide and GSK-J4 are able to block the support provided by YAP* conditioned media, but PR619 is not. Based on this information and knowing the vulnerability of the different cell states, we selected a subset of compounds to see if they could prevent the growth of tumour colonies harbouring multiple resistance mechanisms. PR619 was selected for its ability to target YAP* cells, and SB505124 (TGFβR inhibitor) for its ability to target crosstalk (Fig. 3F), but not YAP* cells, and GSK-J4 and Cilengitide for their ability to target both YAP* cells and crosstalk. Either control Onco- or Onco-NY cells were plated and treated with combinations of PLXPD and the compounds listed above. Figure 7B shows that, as predicted, PLXPD treatment alone was not very effective at eliminating Onco-NY cells. However, combined treatment with PLXPD and GSK-J4 was highly effective. The combination of PLXPD with Cilengitide was also effective, but combination with PR619 or SB505124 was not. These data demonstrate that neither eliminating the YAP* state alone nor blocking inter-subclone crosstalk alone is sufficient to overcome resistance. Instead, effective disease control requires disrupting both the YAP-high cell state and the paracrine signalling network that sustains other resistant cells, a dual function achieved by GSK-J4.

## Discussion

Resistance to cancer therapies has devastating consequences for patients (Russo *et al*., 2024; Sabnis and Bivona, 2019). New technologies have revealed just how heterogeneous advanced malignancies are, and how this heterogeneity is linked to unfavourable outcomes (Qian *et al*., 2017). Autopsy analysis indicates that both multiple genetic resistance mechanisms and multiple cell states are found within a patient, and even within a single lesion (Spain *et al*., 2023; Tirosh *et al*., 2016). Overcoming the challenge of intra-tumour heterogeneity will require either ways to limit heterogeneity or strategies that are effective against multiple different resistant cell states (Patton *et al*., 2021; Tadros *et al*., 2025). To this end, we deliberately set out to generate three different resistant cell states in a model of cutaneous melanoma, addressing a gap in previous studies, which have typically analysed resistance mechanisms separately. One state reflected a recurring genetic change in melanoma resistant to targeted therapy – NRAS* – and the others non-genetic mechanisms – YAP* and SponR. This revealed that the different resistance mechanisms share few common pharmacological vulnerabilities, and that the different resistant states generate an interacting ecosystem with YAP1 active cells providing growth supporting signals for other cells while inhibiting their own growth. Strategies reported to target cells made resistant to targeted therapy via selection, such as HDAC inhibition and GPX4 inhibition leading to ferroptosis (Wang *et al*., 2018; Zhao *et al*., 2025; Maertens *et al*., 2019), were not effective against YAP* cells. We speculate that this may be due to divergent mechanisms of handling redox balance between SponR and YAP*, which is supported by our analysis (Fig. 2B & Fig. 6F).

Numerous studies have shown that melanoma cells can adopt a range of cell states, that can be approximated as a continuum from melanocytic/differentiated to mesenchymal/undifferentiated (Karras *et al*., 2022; Rambow, Marine and Goding, 2019). Moreover, cells are able to transition between states. The A375 model that we use is in neither of these states, but sits in a more intermediate state with drug naïve tumours having low inter-cellular heterogeneity. The transcriptional and epigenetic regulation of melanoma cell states involves many of the same factors involved in the specification of the melanocyte lineage during development, such as MITF, SOX10, YY1, NGFR, and POU3F2 (Brn2)(Seberg, Van Otterloo and Cornell, 2017). Interestingly, several studies have implicated EZH2, which methylates H3K27, as being linked to more aggressive melanoma phenotypes (Zingg *et al*., 2015; Manning, Hooper and Sahai, 2015; Tiffen *et al*., 2016). We identify inhibition of the H3K27 demethylase as an effective strategy to restore sensitivity to targeted therapy. These observations could be reconciled if it is the ability to transition between states that is the key determinant of therapy resistance, and that, even though they have opposing molecular effects, inhibition of EZH2 and JMJD3 both prevent melanoma cells transitioning between states. In the future, it will be important to map the relationship between melanoma states and H3K27 methylation and to determine the extrinsic factors driving cells between different states in tumours. Previous studies have indicated that αMSH can drive melanoma cells into melanocytic states while TGFβ promotes more mesenchymal states (Manning, Hooper and Sahai, 2015; Nishimura *et al*., 2010). Consistent with the latter observation, we find that blocking TGFβR signalling reduces the ability of YAP* cells to promote therapy resistance in control cells.

Intra-tumour heterogeneity, long observed in patient tissues, suggests targeting inter-subclone signalling may have broad utility across cancers. Paracrine IL11 signalling has been shown to promote the maintenance of intra-tumour heterogeneity in breast cancer (Marusyk *et al*., 2014), and emergent synergy between MYC high and low and WNT high and low states has been reported in genetically engineered breast and lung cancer models (Kreuzaler *et al*., 2019; Tammela *et al*., 2017; Wang *et al*., 2015). Thus, when considering the microenvironment of cancer cells, it is important to consider paracrine signals that might be emanating from nearby cancer cells in different states, not just signals from non-cancerous cells. However, experimental systems that enable control of tumour heterogeneity remain rare. Here we adapt the Brainbow lineage labelling cassette to enable the induction of heterogeneity in established tumours. The application of our system and of combined Cre-recombinase and Flippase genetically engineered models should enable further experimental dissection of how intra-tumour heterogeneity impacts cancer progression and therapy responses. In the longer term, improved understanding of the mechanisms sustaining intra-tumour heterogeneity, such as inter-subclone communication, should enable more effective therapeutic strategies to treat tumours composed of heterogeneous cell states.

## Material and Methods

### Generation of targeted therapy resistant cell lines

A375P (A375) cells and their derivatives generated were cultured in DMEM media with 10% FBS, 100 U/ml Penicillin and 100 µg/ml Streptomycin at 37°C and 5% CO_2_.

The A375 human BRAF^V600E^-mutant melanoma cells were infected either with an empty retroviral vector (pBabe) or an empty lentiviral vector (PCSII-fluorophore-CAAX-IRES-Hygro) to generate control cell lines (A375 CTR). A375 NRAS* cells were generated by infection with either a pBabe or PCSII vector expressing the mutated NRAS^Q61K^ while A375 YAP* cells were infected with either a pBabe or PCSII expressing the constitutively active form YAP1-5SA, in which five serines (S61, S109, S127, S164, S381) have been exchanged for alanine and thus cannot be phosphorylated. Long-term culture of A375 CTR cells in BRAF+MEKi (1 μM PLX-4720 + 1 µM PD184352 = 1 µM PLXPD) for 19 weeks resulted in a spontaneous resistant cell line (A375 SponR). A375 SponR cells were maintained in culture hereafter with 0.5µM PLXPD. A375 cells containing either the empty Oncobow cassette or Oncobow incorporating NRAS^Q61K^ and YAP1^5SA^ were generated through transposase-mediated integration of the PiggyBac vector containing them followed by selection using puromycin (2µg/ml), single cell cloning, and testing for low recombination baseline and high levels of induced recombination.

### Focus assays

Cells were seeded in 24-well plates (A375 CTR, NRAS* and YAP* cell lines: 7500 cells/well, A375 SponR cells: 12500 cells/well, HaCaT cells: 12500 cells/well, 4434 CTR, NRAS*, YAP* and SponR cell lines: 7500 cells/well; H1975 CTR, KRAS^G12C^ and KRAS^G12D^ cell lines: 7500 cells/well). After 24 hours, respective drug treatment was added. Media and drugs were changed every 2 to 3 days and the assay was stopped at day 7 or 8 after drug administration. At the end point, the media was removed, cells washed with PBS and fixed with 4% paraformaldehyde for 10 minutes at room temperature. Subsequently, the fixative was removed, cells washed in PBS and stored with PBS in the fridge until further processing for immunofluorescence, imaging or crystal violet staining.

### Focus assays for Oncobow cells

For long-term focus assays, 2000 A375 Onco or A375 Onco-NY cells were seeded per well in 24-well plates (ibidi µ-Plate 24-well, 82426). After 5h, cells were treated with ethanol (non-recombination control) or 20nM 4-hydroxytamoxifen (4-OHT). 4-OHT or ethanol was replaced after 18h with PBS washes followed by normal media. After 2 days, drugs were added, refreshed every 3-4 days, and the assay concluded 13 days post-drug addition by PBS washing and fixation with 4% PFA for 10 min at room temperature. Fixed cells were washed, stored in PBS at 4°C, and processed for immunofluorescence, imaging, or crystal violet staining.

### Crystal violet staining

Fixed cells were incubated with 0.05% crystal violet solution (in ddH_2_O) for 30min. Crystal violet solution was removed and cells washed twice with tap water before plates were left to dry and scanned using an EPSON Perfection V700 Photo scanner.

### Small Molecule Inhibitor Screen

A375 and HaCaT cells were passaged as described above and seeded in 96-well plates with 100 µL media per well (A375 cell lines: 3000 cells/well, HaCaT cells: 4000 cells/well. After 24 hours, compounds (Table 1) were added and cells incubated for 72 hours with drug. Media of death control wells was removed, 10% Triton X-100 added and incubated for one hour. Then, 100 µL CellTiter-Glo^®^ reagent was added, incubated for 10 minutes at room temperature and luminescence was measured using the TECAN infinite^®^ M1000. The data were analysed using R 4.0.5 (R Core Team, 2021).

**Table 1.**
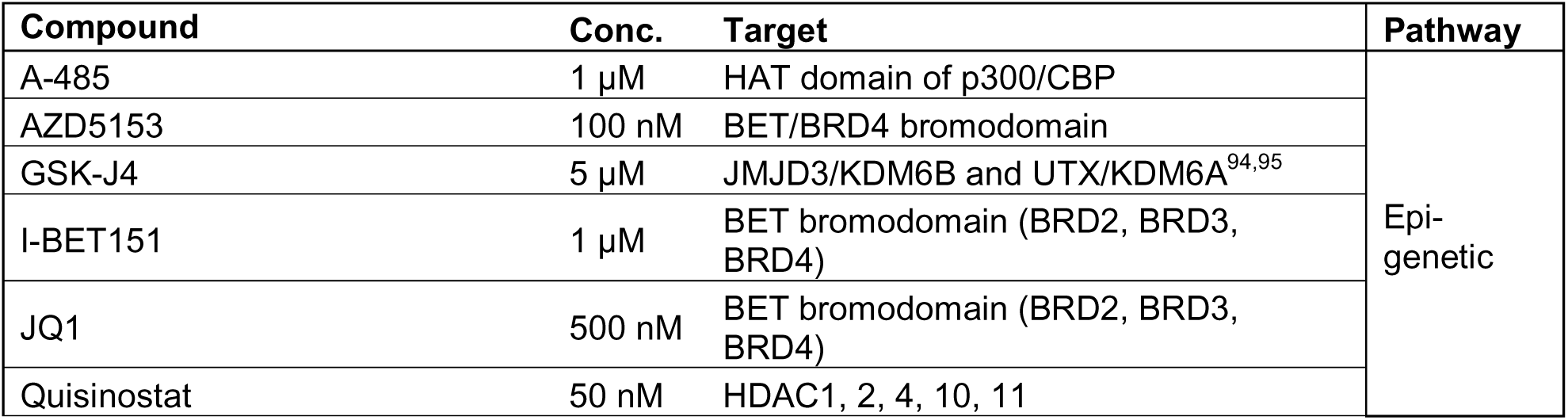
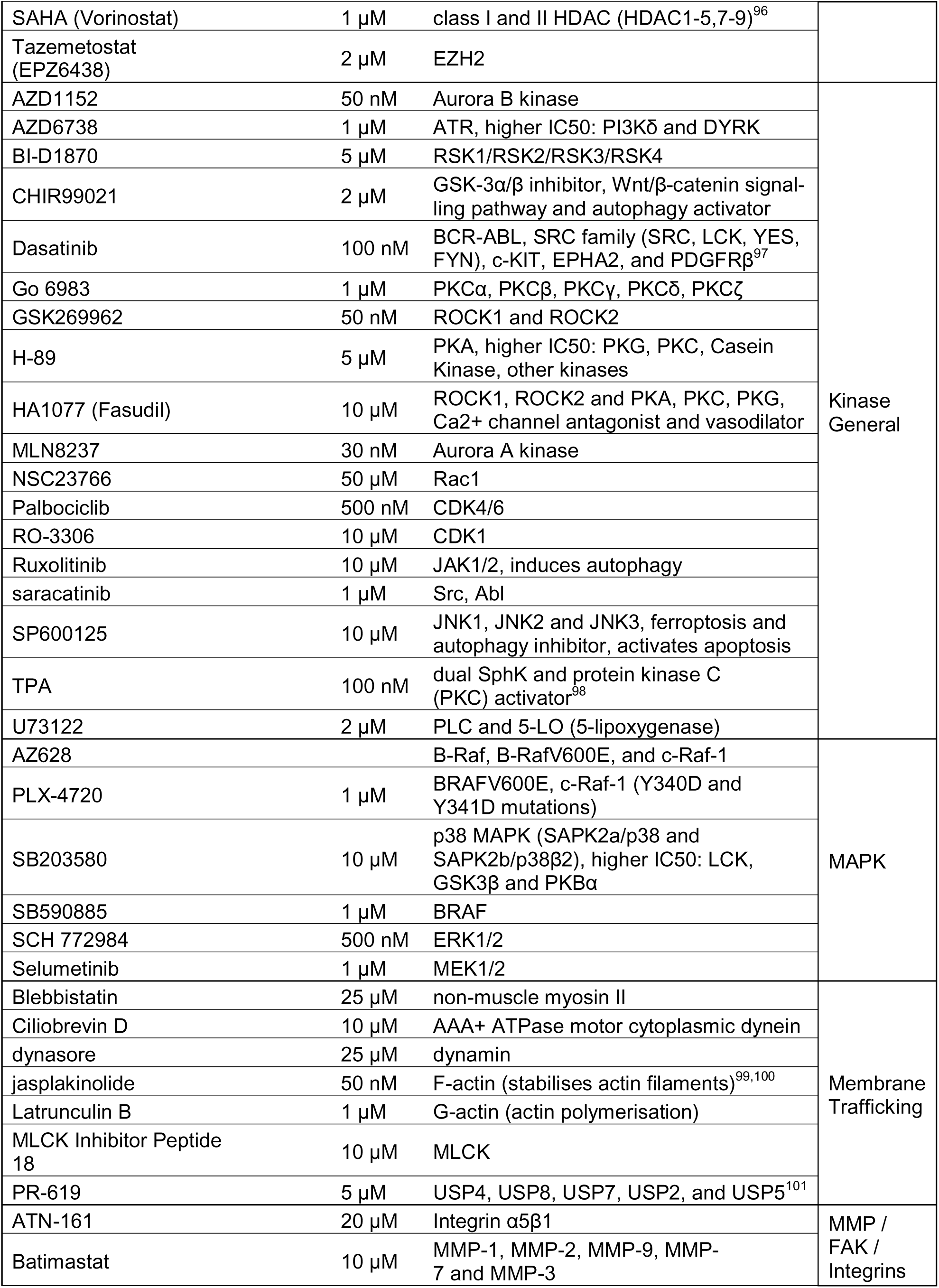
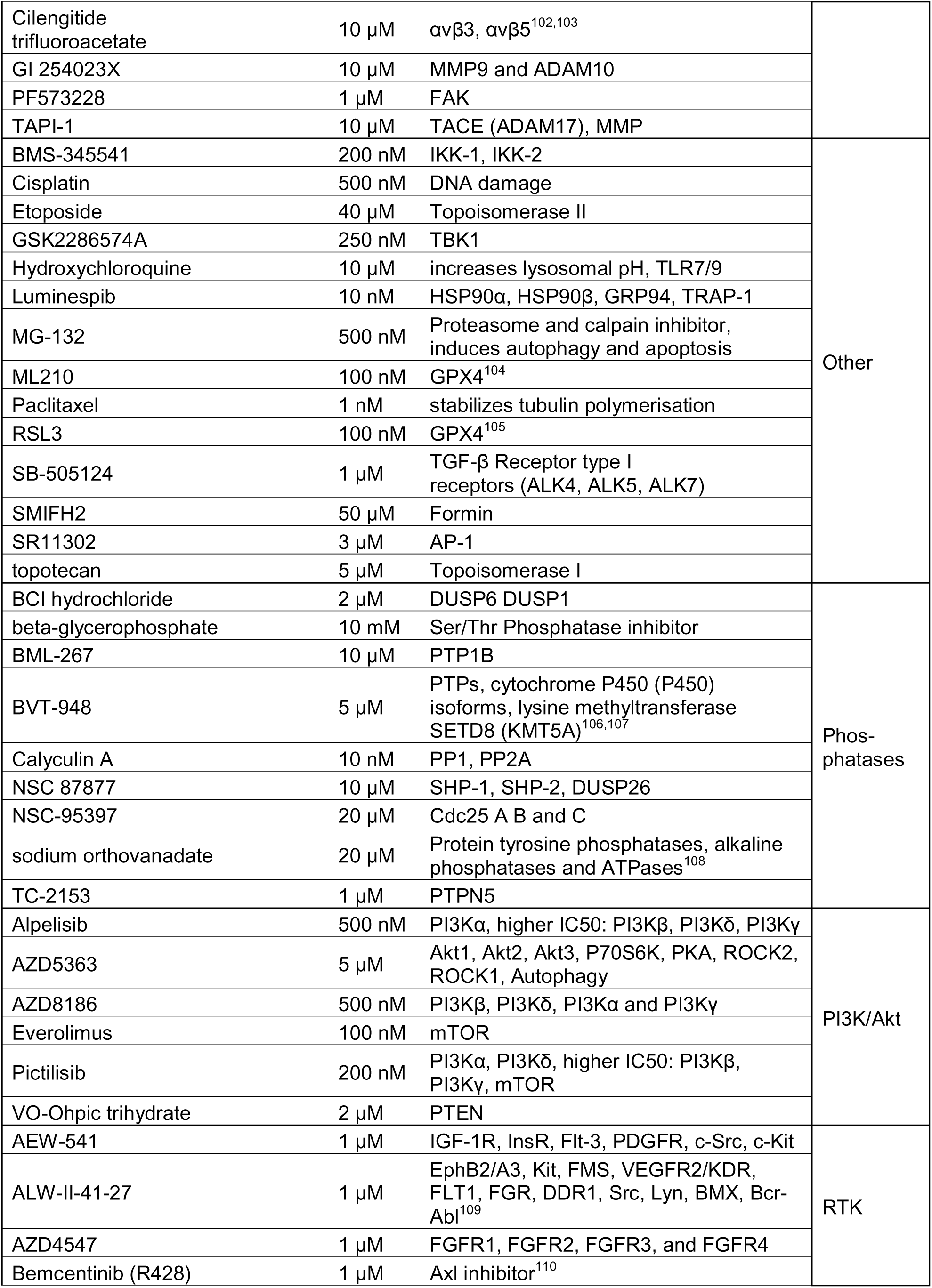
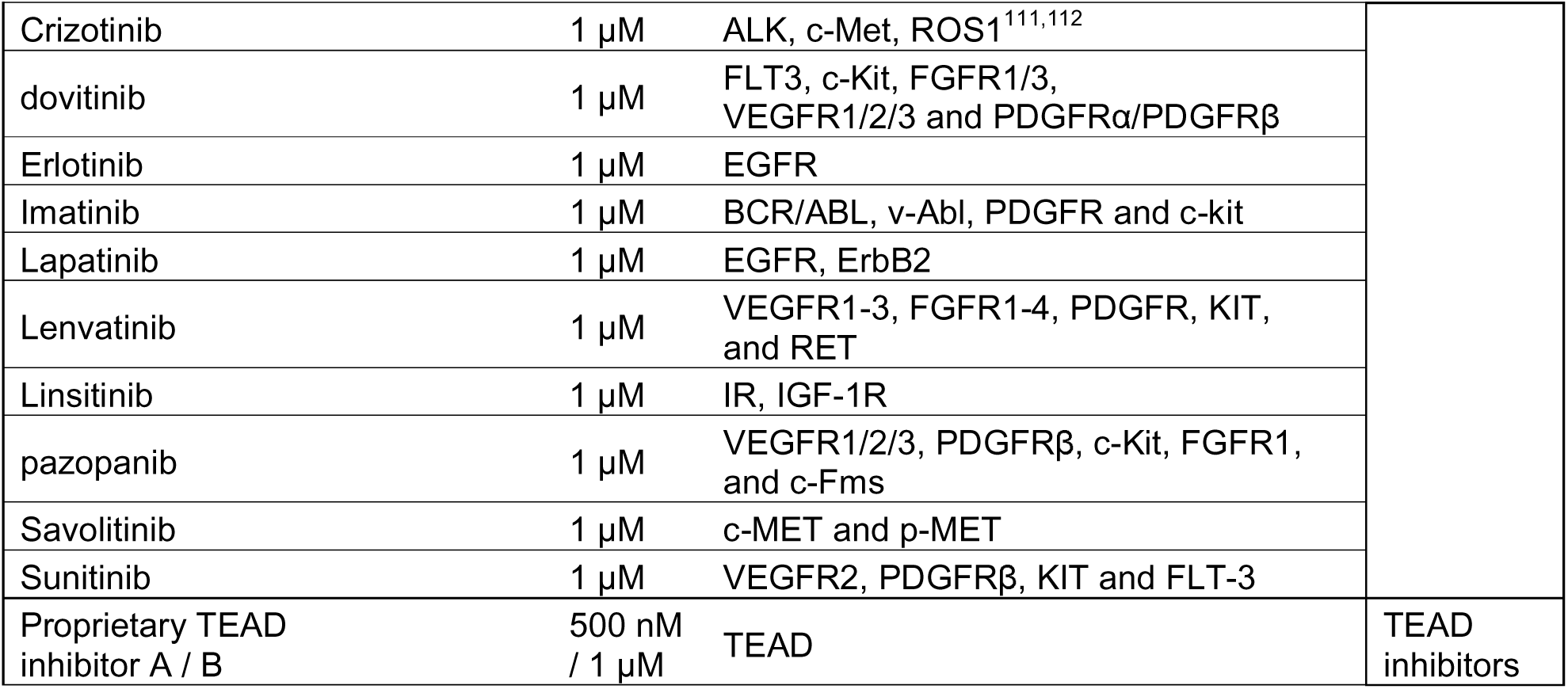
Compounds used in the pharmacological lethality screen. Compounds with used working concentration, target(s) and related pathway.

### Incucyte growth analysis

To measure cell growth, 7500 cells/ 24-well were seeded in triplicate for the A375 CTR, NRAS*, YAP* and SponR cell lines. For experiments including PLXPD treatment, media was removed 24 hours after seeding and media with either 0.02% DMSO or 1 μM PLXPD was added and cell growth was monitored in the Incucyte for 5/6 days.

### In vivo mouse experiments

For in vivo subcutaneous tumour growth experiments for A375 CTR, NRAS*, YAP* and SponR cells 1x10^6^ cells were injected in 100µL PBS/Matrigel (1:2) in the flank of NOD-SCID mice. When tumours reached 40-60 mm^3^ in volume (approximately 13 days after injections) mice were randomised and treatment commenced with DMSO or the combination of PLX4720 (25mg/kg) and Trametinib/ GSK1120212 (1mg/kg) diluted in water by oral gavage, daily/3 times per week. For A375 Onco or Onco-NY cells, 5 x 10^5^ cells cells were injected in 100µL PBS/Matrigel (1:2) in the flank of NSG mice. Once tumours were palpable (2-3mm), mice received two doses of 100µL 4-hydroxytamoxifen (20mg/kg in saline) intraperitoneal injection on two consecutive days. When tumours reached 40-60 mm^3^ in volume (typically 20-22 days after injection) mice were randomised and treatment commenced with DMSO or the combination of PLX4720 (25mg/kg) and Trametinib/ GSK1120212 (1mg/kg) diluted in water by oral gavage, daily/3 times per week.

### Protein extraction, quantification and Western Blot analysis

A375P cells (CTR, NRAS*, YAP*) were seeded at 1×10□ cells/well, and SponR cells at 2×10□ cells/well in a 6-well plate. After 24 h, cells were treated with DMSO or 1 µM PLX4720 and 1 µM PD184352 for 24 h. Cells were washed three times with cold PBS and lysed in 150 µL RIPA buffer (50 mM Tris-HCl, 150 mM NaCl, 1 mM EDTA, 1% Triton X-100, 1% sodium deoxycholate, 0.1% SDS) supplemented with protease and phosphatase inhibitors (PhosSTOP tablet Roche #04906837001, cOmplete EDTA-free Roche #11873580001, 50 mM NaF). Lysis was performed directly in plates using a cell scraper, followed by 10 min on ice and centrifugation (16,000 g, 30 min, 4°C). Protein quantification was performed via the bicinchoninic acid (BCA) method (ThermoFisher #23225). For Western blotting, protein was loaded onto a 4–15% Mini-PROTEAN TGX gel (Biorad #4561084) and transferred to a Trans-Blot Turbo Mini 0.2 µm PVDF membrane (Biorad #1704156). The membrane was blocked in 5% BSA or milk in TBST, incubated with primary antibodies (overnight at 4°C or 1 h at room temperature), followed by HRP-conjugated secondary antibodies (ThermoFisher). After washing, detection was performed using Luminata Classico Western HRP substrate (Millipore #WBLUR0100, #WBLUF0100). Antibody details are in Table S4, and original blots are provided as source data.

### ROS assay

A375P cells and their derivatives were seeded in a 96 well glass-bottom Ibidi plate at 20,000 per well for the SponR cells or 10,000 for control, NRAS and YAP and incubated for 24 hours to adhere. Drug treatment was added for 72 hours. Following treatment, media was removed and replaced with fresh media containing 5 uM cellROX green reagent (Thermo Fisher C10444) and Hoechst. Plate was incubated for 30 minutes at 37C, washed 3 times with PBS and imaged with a Zeiss LSM980. The analysis was done in ImageJ by generating a nuclear ROI using the Hoechst channel, and measuring the mean intensity of the cellROX channel for each cell.

### RNA sequencing

1 x 10^6^ A375P cells and their derivatives were seeded per well (6 well plates) and incubated for 48 hours. Then, plates were washed once with PBS, buffer RLT with 1% β- mercaptoethanol was added and cells were detached by cell scrapers. RNA was isolated with the RNeasy® Mini Kit and RNase-free DNase Set according to the manufacturer’s protocol, the final elution was performed with 30 μl RNase-free H2O. The RNA concentration was measured using the NanoDrop® ND-1000 Spectrophotometer.

### Immunofluorescence and immunohistochemistry

A375 cells were seeded in a 96 well glass-bottom Ibidi plate at 10,000 per well for the SponR cells or 2500 per well for control, NRAS and YAP. Plates were incubated for 24 hours to adhere, then media was aspirated and replaced with fresh media containing drug treatments for 24 hours. Cells were fixed in 4% PFA for 15 minutes at RT, permeabilized in 0.2% Triton X-100 for 15 mins at RT and blocked in 5% BSA overnight at 4C. Primary antibodies integrin β3 (Santa Cruz, sc13579, 1:00), YAP1 (Cell Signaling D8HX1), or paxillin (BD Transduction Laboratories, 610051, 1:100) were incubated in 0.05% Triton X-100 and 1.5% BSA overnight at 4C, then washed three times in 0.01% Tween-20 in PBS. The secondary antibody donkey anti-mouse Alexa 647 (Invitrogen, A31571, 1:1000) with DAPI (Sigma Aldrich, D-9542,1:1000) and phalloidin (Sigma Aldrich, P5282, 1:500) was incubated in 0.05% Triton X-100 and 1.5% BSA for 1 hour at RT, then washed three times in 0.01% Tween-20 in PBS. Images were acquired with a Zeiss LSM980 confocal microscope. Immunohistochemistry of formalin-fixed paraffin-embedded tissue was performed using standard methods and the following antibodies: YAP1 (Cell Signaling D8HX1), Collagen I (Abcam Ab215969), and GFP (Thermo Fisher A-6455).

### Highly Multiplexed Immunostaining and analysis

Multiplexed immunofluorescence was performed on a subset of diagnostic blocks from the melanoma TRACERx clinical study, using the PhenoCycler-Fusion system (Akoya Biosciences) as previously described in Yoon, Novo, et al in revision Cell Systems (Yoon *et al*., 2024). Briefly, 5 µm FFPE sections underwent heat-induced antigen retrieval (TE buffer, pH 9.0) followed by blocking and a 3 h incubation with an oligonucleotide-conjugated antibody cocktail (detailed below). Following post-staining fixation, sections were imaged via automated cycles of complementary fluorophore-conjugated reporter addition and removal. Specific antibody dilutions and exposure settings are detailed in Table 2. QuPath was used to segment nuclei and their associated cytoplasm. Melanoma nuclei were determined by SOX10 staining >5. Principal Component Analysis was performed using GraphPad Prism.

| Target | Antibody clone | Vendor & Catalogue Number |
| --- | --- | --- |
| aSMA | 1A4 | Novus Bio NBP2-34522 |
| Beta III Tubulin | EP1569Y | Abcam ab221935 |
| CAIX | EPR4151 (2) | Abcam ab108351 |
| CD103 | EPR4166 (2) | Abcam Ab271889 |
| CD11c | 118/A5 | Akoya 4550135 |
| CD20 | L26 | Akoya 4450018 |
| CD206 | E2L9N | CST 49243SF |
| CD31 | EP3095 | Akoya 4450017 |
| CD3e | EP449E | Akoya 4550119 |
| CD4 | EPR6855 | Akoya 4550112 |
| CD45 | D9M81 | Akoya 4550121 |
| CD68 | KP1 | Akoya 4550113 |
| CD79a | D1X5C | Akoya 4450078 |
| CD8 | C8/144B | Akoya 4250012 |
| Cleaved Caspase 3 | 5A1E | CST 94530SF |
| Collagen I | EPR7785 | Abcam Ab215969 |
| Collagen IV | EPR20966 | Akoya 4550122 |
| E-cadherin | 4A2C7 | Akoya 4250021 |
| FAP | EPR20021 | Abcam Ab271976 |
| Fibronectin | E5H6X | CST 44800SF |
| FoxP3 | 236A/E7 | Abcam Ab96048 |
| GP100 | HMB45 | Akoya 4550091 |
| GZMB | EPR20129-217 | Abcam Ab219803 |
| HLA-ABC | EPR22172 | Abcam Ab239788 |
| Integrin Beta III | ERP17507 | Abcam ab240214 |
| Ki67 | B56 | Akoya 4250019 |
| Melan A | A103 + M2-7C10 + M2-9E3 | Abcam ab212366 |
| MPO | E1E7I | CST 88757SF |
| Pan-Cytokeratin | AE-1/AE-3 | Akoya 4450020 |
| PD-1 | D4W2J | Akoya 4550038 |
| PD-L1 | 73-10 | Akoya 4550128 |
| Podoplanin | NC-08 | Akoya 4250004 |
| Sox10 | BSB-62 | Biolegend 847202 |
| Tenascin C | 4C8MS | Novus Bio NB110-68136 |
| THBS1 | EPR2297-54 | Abcam ab267397 |
| Vimentin | D21H3 | CST 46173SF |
| Yap | D8HIX | CST 53921SF |

### Migration assay

7500 A375P cells or derivatives were seeded in 24 well plate (Ibidi 82426) coated with a mixture of 150µl of 2mg/ml Matrigel (Corning #354234) and 4mg/ml Collagen I (Corning #354249) gel. After 24 hours 1µM PLX4720 (Selleck Chem S1152) and 1µM PD184352 (Selleck Chem S1020) or DMSO were added. Imaging using phase contrast microscopy was commenced after 48 hours, with 1µM SiR-DNA added (Cytoskeleton Inc. cy-sc007) 1hr prior to imaging. Three positions were imaged per well for 20hrs with a 10min frame rate.

### Determination of differential ligand expression between resistant states

To determine differential ligand expression between resistant states, we downloaded the complete list of ligands from the NicheNet R package (v2.1.5) and used the variance-stabilized transformation (VST) normalized counts from the transcriptomic dataset. We then compared the expression of each ligand in the following samples: CTR_DMSO, NRAS*_DMSO, YAP*_DMSO and SponR_DMSO. To identify the number of ligands specifically expressed in each sample, we performed a fold-change comparison of the VST-normalized count for each sample against the others. A ligand was considered enriched in a specific sample if it showed at least a 1.3-fold increase compared to each of the other samples.

### Deriving signature linked to BRAFi + MEKi resistance

To derive a gene signature associated with BRAFi + MEKi resistance, we used the VST normalized counts from the transcriptomic dataset. We selected genes that were upregulated in NRAS*_PD, YAP*_PD, SponR_PD, and compared to CTR_PD, with a fold-change increase of at least 1.3.

### NicheNet analysis to determine which differentially expressed ligands can drive gene expression linked to resistance

Intercellular communication was assessed using an adapted version of the NicheNet R package (v2.1.5) for bulk RNA-seq analysis, following the vignette available at <u>this link</u>. The analysis was conducted using the sending_YAP_DMSO_ligands_2.R script.

First, the human ligand-receptor network was loaded from the NicheNet Zenodo repository, followed by the variance-stabilized (VST) normalized counts dataset. The list of ligands expressed in the YAP*_DMSO sample was obtained as described in the section “Determination of differential ligand expression between resistant states.” Specifically, ligands were selected based on their expression in YAP*_DMSO compared to CTR_DMSO and NRAS*_DMSO. The list of receptors in the receiving cells was determined by selecting genes with expression levels exceeding 11 normalized counts. These expressed ligand and receptor lists were then filtered to include only those pairs that putatively interact according to the NicheNet database.

For the gene set of interest, we used the BRAFi + MEKi resistance signature defined in the previous section. Intercellular communication analysis was performed using these inputs to predict ligand activities (predict_ligand_activities) and infer target genes and receptors of top-ranked ligands (get_weighted_ligand_target_links). Data visualization was carried out using NicheNet visualization functions and GraphPad Prism, and all comparisons followed the same analytical steps. Full details can be found in the aforementioned script.

To compare ligand activity across different samples, we used the area under the precision-recall curve (AUPR) corrected value and applied z-score normalization for each sample. If a ligand was not predicted to have an effect, the minimum value was assigned for visualization in the corresponding heatmap.

### Conditioned media experiments

To generate conditioned media, 4 × 10□ A375 CTR or YAP* cells were seeded per T175 flask and cultured for 4 days. The media was then collected and filtered through a 0.45 µm filter. To prepare the final media, one-third fresh DMEM (supplemented with 10% FBS and 1% Pen-Strep) was mixed with two-thirds of the conditioned media. This mixture was subsequently frozen at -80°C until use.

For receiving cells, 6 × 10³ CTR, NRAS*, YAP*, or SponR cells were plated per well in a 24- well ibidi µ-Plate. One day after seeding, cells were treated with either 0.02% DMSO or 1 µM PLXPD. Drug treatments were refreshed every 3 days, and the assay was terminated 7 days after initial drug addition. After treatment, cells were gently washed once with PBS and fixed using 4% paraformaldehyde for 10 minutes. Following a PBS wash, plates were stored at 4°C until staining. For phalloidin staining, cells were permeabilized with 0.1% Triton X-100 for 10 minutes, washed with PBS, and then incubated with Phalloidin-Atto 633 (1:500) and DAPI (1:1000) in a solution of 0.1% Tween-20 and 1% BSA in PBS for 30 minutes to 1 hour. Plates were subsequently washed twice with 0.1% Tween-20 in PBS, once with PBS, and stored in PBS at 4°C until imaging with the Olympus FV3000.

To test if compounds are able to hinder the supportive signal emanating from YAP* cells to CTR compounds were added in combination with 1µM PLXPD 1 day (concentrations see table x) after seeding 6 × 10³ A375 CTR cells per well in a 24-well ibidi µ-Plate. Drug treatments were refreshed every 3 days, and the assay was terminated 10 days after initial drug addition and fixation and phalloidin staining performed as described above.

| Compound | Target | Concentration |
| --- | --- | --- |
| PR619 | DUBs | 2μM |
| Cilengitide | Integrin b3 | 5μM |
| GSK-J4 | JMJD3/KDM6B and UTX/KDM6A | 1μM |
| Dasatinib | BCR-ABL, SRC family (SRC, LCK, YES, FYN), c-KIT, EPHA2, and PDGFRβ | 100nM |
| GSK269962 | ROCK1, ROCK2 | 50nM |
| PF573228 | FAK | 1μM |
| SB-505124 | TGFR1, ALK4, ALK5, ALK7 | 1μM |
| AZD4547 | FGFR1-4 | 1μM |
| ACT001 | PAI-1 | 10μM |
| Erlotinib | EGFR | 1μM |

### Agent based mathematical modelling

An agent-based model was developed to describe cell proliferation, pushing and pulling forces between adjacent cells, cell random motility, and cell death. Briefly, cell proliferation is controlled both intrinsically, through sampling at cell birth a doubling time *T^double^* from a distribution *X* ∼Erlang(*k*, Λ) with its shape and rate Λ derived from *T^mean^* = *k*⁄Λ and 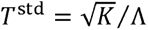 and extrinsically, in response to the compressive forces from surrounding cells. Cell random motility is modelled by introducing a stochastic noise to its coordinate vector. Components of this noise, 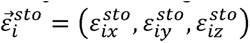, are represented by normally distributed random variables. The *x* component is expressed as 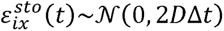, where is the diffusion coefficient and Δ is a time step. Cell death is modelled under therapy, with a probability Computationally, a random number is generated is generated between [0, 1). If *j < P^death^*, a cell under consideration is eliminated from the simulation. In this study, therapy is administered continuously once a population reaches 10000 cells.

Considering experimental observations of phenotypic differences between cell states in this study, several parameters for cell traits were chosen differently in a cell-state-specific manner, as shown in the table below. In addition, actual cell traits related proliferation and therapy-induced death are further modulated during a simulation according to inter-clone interactions (see below).

|  | CTR, CTR-labelled | NRAS | YAP |
| --- | --- | --- | --- |
| $T^{\text{mean}}$ | 20 hour | 40 hour | 20 hour |
| $C$ | $5 \times 10^4$ | $5 \times 10^4$ | $4 \times 10^5$ |
| $D$<br>($1 \times 10^2 \mu\text{m}^2 \text{h}^{-1}$) | 0.28 | 0.28 | 2.22 |
| $p^{\text{death}}$ | 0.0004 | 0.004 | 0.004 |

\*Note: Motility factor $C$ (and relatedly diffusion coefficient $D$ ) for CTR and NRAS shown in the table reflects value used for representative simulations. Further simulations with varying levels of motility factors were performed, reflecting varying extents of migratory advantage of YAP cells, and respective outputs reported in this study.

#### Modelling of inter-clone interactions

Considering experimental observations, the model assumes a self-inhibitory effect on proliferation for YAP cells. The sampled intrinsic doubling time of a YAP cell born at a certain time *t_1_* is scaled by a factor 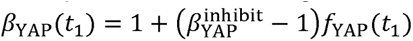, where 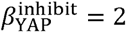 in this study and *f^YAP^ (t)* is the fraction of YAP cells out of the total cell population at time *t_1_*.Namely,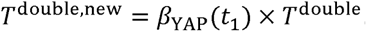 In addition, the model assumes a boosting effect on proliferation of CTR or NRAS cells by YAP cells. The sampled intrinsic doubling time of a CTR or NRAS cell born at a certain *t_2_* time is scaled by a factor 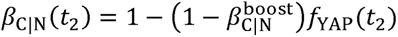 where 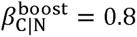 for representative simulations in this study and *f_YAP_ (t_2_)* is the fraction of YAP cells out of the total cell population at time. Further simulations with varying levels of 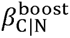 within a range of [0.5, 1] were performed and respective outputs reported in this study. Namely,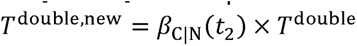.

Furthermore, the model assumes a boosting effect on survival under therapy of CTR or NRAS cells by YAP cells. The death probability of a CTR or NRAS cell under therapy at a certain time *t_3_* is scaled by a factor 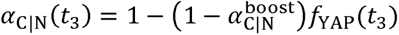, where 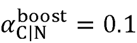 in this study and *f^YAP^(t_3_)*is the fraction of YAP cells out of the total cell population at time *t_3_*. Namely,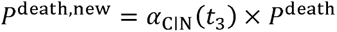.

#### Simulation with stochastic cell state transitions

In a set of *in silico* experiments, a simulation starts with a single CTR cell. Every 1 hour of simulation time, stochastic cell state transitions take place governed by a probability *p*^transition^. Computationally, the following procedure is iterated over all CTR cells. A random number *k* is generated between [0, 1). If *K < p*^transition^, the CTR cell transitions to YAP, NRAS, or CTR-labelled (with no phenotypic difference compared to CTR) state with equal probability *p*^transition^ 0.00003 for representative simulations in this study. Further simulations with lower and higher levels of *p*^transition^ were performed and respective outputs reported in the study.

#### Simulation with induced cell state transitions

In a set of *in silico* experiments, a simulation starts with a single CTR cell. At a predefined time *T* = 216 hour, a random fraction of cells *F*^induce^=0.2 are selected to undergo state transition to YAP, NRAS, or CTR-labelled (with no phenotypic difference compared to CTR) state with equal probability.

#### Visualisation

Snapshots of cell populations from simulations are generated by visualising cell positions using scatter plots. CTR, YAP, NRAS, and CTR-labelled cells are visualised in grey, yellow, turquoise, and red, respectively.

## Acknowledgements and Disclosures

The authors thank all members of the Sahai lab for their insightful comments on the manuscript. **K.S.** received funding from the Deutsche Forschungsgemeinschaft (DFG) and the European Research Council (ERC Advanced Grant CAN_ORGANISE, Grant agreement number 101019366). **G.G.** received funding from the European Research Council (ERC Advanced Grant CAN_ORGANISE, Grant agreement number 101019366). **A.C.** received funding from the European Research Council (ERC Advanced Grant CAN_ORGANISE, Grant agreement number 101019366). **S.B., S.H., S.M.,** and **E.S.** is supported by the Francis Crick Institute, which receives its core funding from Cancer Research UK (CC2040), the UK Medical Research Council (CC2040), and the Wellcome Trust (CC2040) and the European Research Council (ERC Advanced Grant CAN_ORGANISE, Grant agreement number 101019366).

**E.S.** reports grants from Novartis, Merck Sharp Dohme, AstraZeneca and personal fees from Phenomic.

## Author contributions

K.S. and E.S. devised the study. K.S. performed the majority of experiments. S.B. helped with ‘Oncobow’ experiments. S.H. helped generate the ‘Oncobow’ cells and performed many of the pre-clinical experiments. V.H. performed the in vitro drug screening. S.M., A.C., and L.T. assisted with in vitro experiments. G.G. and S.S. performed bioinformatic analysis. A.B. provided pathology advice. X.F. performed the computational simulations. B.S. and S.T. provided human melanoma samples and advice.

## Supplementary Movies

Movie 1 corresponds to Figure 4C. The simulated growth of a tumour containing with a slow rate of spontaneous transitions to different states continually occurring and no treatment. Yellow indicates YAP high state, green indicates Ras mutant state. red indicates no change in state.

Movie 2 corresponds to Figure 4C. The simulated growth of a tumour containing with a slow rate of spontaneous transitions to different states continually occurring and treatment initiated at day 12. Yellow indicates YAP high state, green indicates Ras mutant state, red indicates no change in state.

Movie 3 corresponds to Figure 5D. The simulated growth of a tumour containing the empty Onco-cassette with clonal labelling triggered at day 10 and no treatment. All colours indicate no change in cell state.

Movie 4 corresponds to Figure 5D. The simulated growth of a tumour containing the empty Onco-cassette with clonal labelling triggered at day 10 and treatment initiated at day 12. All colours indicate no change in cell state.

Movie 5 corresponds to Figure 5D. The simulated growth of a tumour containing the NY Onco-cassette with clonal labelling triggered at day 10 and no treatment. Yellow indicates YAP high state, green indicates Ras mutant state, red indicates no change in state.

Movie 6 corresponds to Figure 5D. The simulated growth of a tumour containing the NY Onco-cassette with clonal labelling triggered at day 10 and treatment initiated at day 12. Yellow indicates YAP high state, green indicates Ras mutant state, red indicates no change in state.

---

**Supplementary Figure 1.**
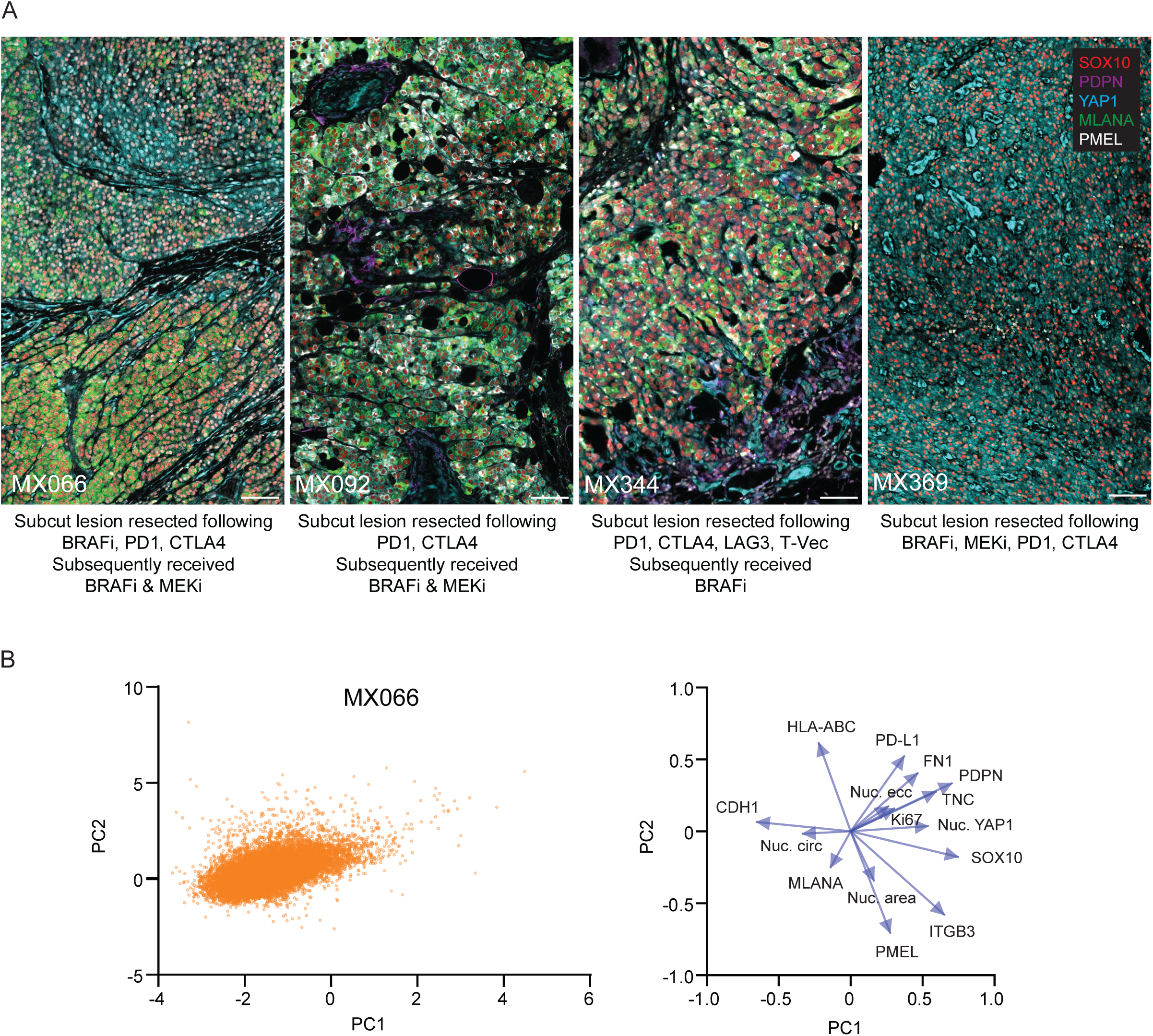
Heterogeneity in cell state of therapy resistance melanoma. A. Images show immunostaining using Phenocycler platform of 4 different sub- cutaneous melanoma lesions treated as indicated. Staining highlights SOX10 (red), YAP1 (cyan), PDPN (magenta), MLANA (green), PMEL (white). Scale bar is 100µm. B. Left-hand plot shows PCA of cells from sample MX066. Right-hand plot shows the PCA loadings.

**Supplementary Figure 2.**
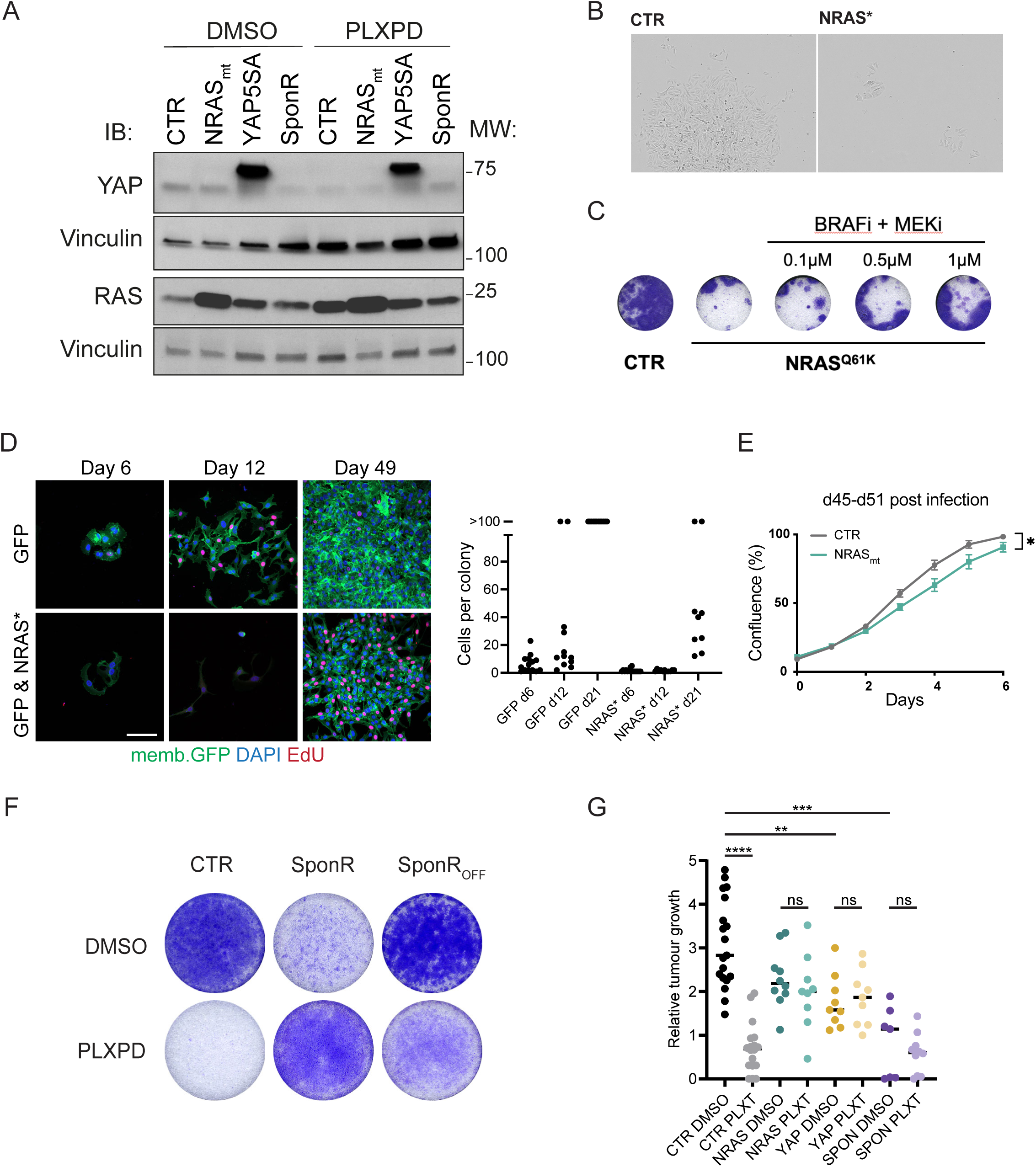
Establishment of three different BRAF/MEKi resistant phenotypes in a single model. A. Western blot shows the expression of endogenous RAS isoforms and YAP1 and ectopically expressed forms in NRAS* and YAP* cells, with vinculin used as a loading control. Molecular weight markers are shown on the right. B. Images show the typical difference in the size of colonies following infection with control virus, pBabe puro, or NRAS^Q61K^ virus. C. Images show number of colonies formed per well after 8 days of culture with or without the indicated levels of BRAFi (PLX4720) and MEKi (PD184352). D. Images and plot show colonies and colony sizes, respectively, of A375 cells 6, 12, and 49 days after transduction with control GFP-CAAX or NRAS^Q61K^ GFP-CAAX virus. In images DAPI is blue, EdU is red, and GFP is green. Scale bar is 100µm. E. Plot shows the growth rate of control and NRAS^Q61K^ expressing cells 45-51 days post infection. F. Crystal violet staining is shown to indicate the growth of the indicated cells (CTR, SponR, and SponR_OFF_, which are SponR cells transferred back into PLXPD-free media for at least two weeks) 8 days after seeding. G. Plot shows tumour growth between days 13 and 21/22 for the indicated cells with or without PLX and Trametinib treatment of the mice.

**Supplementary Figure 3.**
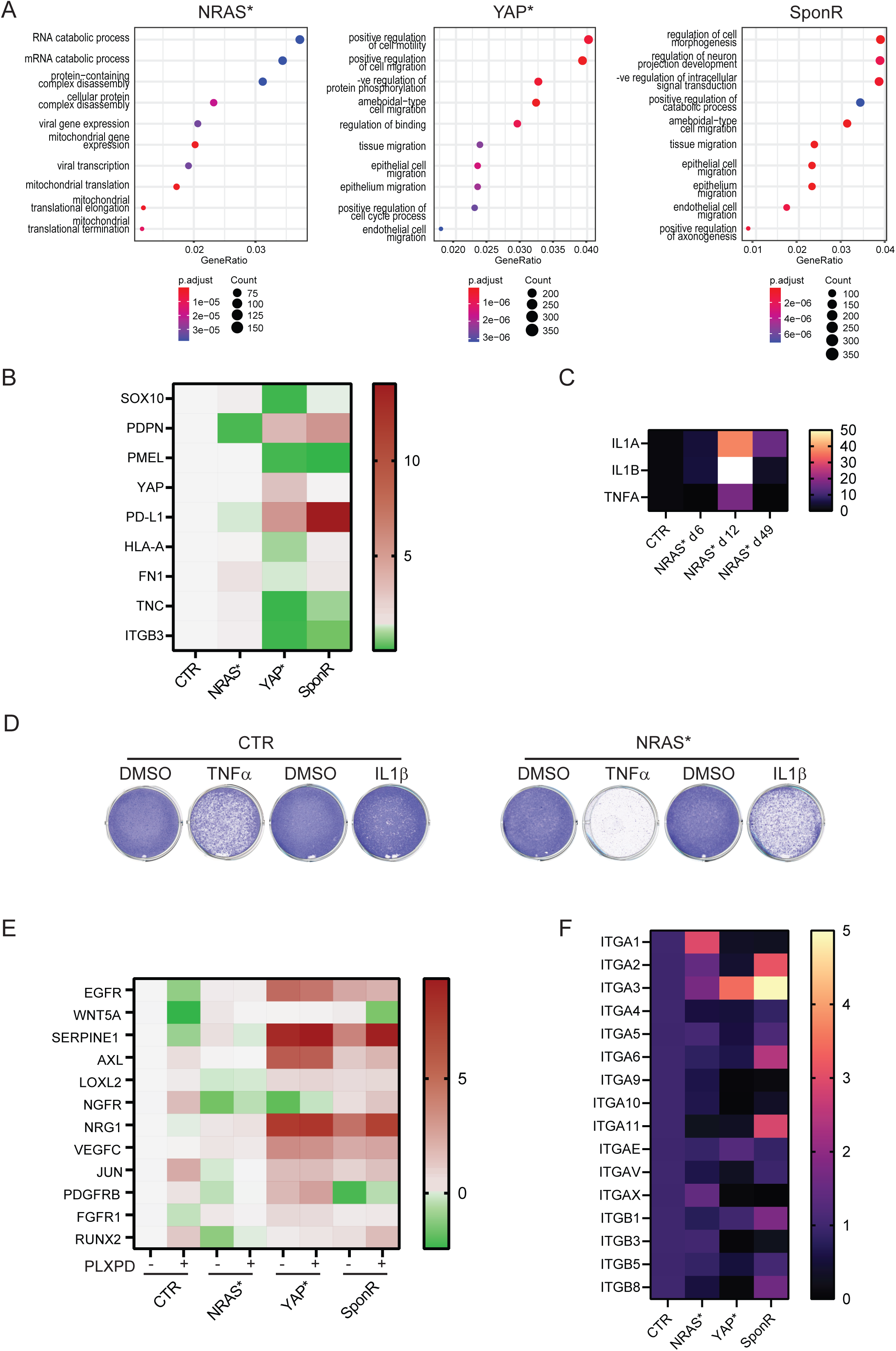
Characterization of three different BRAF/MEKi resistant phenotypes. A. Plots show the top ten most up-regulated transcriptional programmes (as defined by Gene Ontology terms) in NRAS*, YAP*, and SponR cells. RNAseq – more details B. Heatmap shows the expression relative to CTR cells of the indicated genes in NRAS*, YAP*, and SponR cells. C. Heatmap shows the expression of IL1A, IL1B, and TNF in cells transduced with NRASQ61K compared to empty vector controls at the indicated timepoints. D. The effect of TNFα on the growth of CTR and NRAS* cells is shown – visualised by crystal violet staining 8 days E. Panel shows the changes in expression of genes linked to BRAFi + MEKi resistance by Schaffer et al in NRAS*, YAP*, and SponR cells. F. Panel shows the changes in integrin genes linked to BRAFi + MEKi resistance by Schaffer et al in NRAS*, YAP*, and SponR cells.

**Supplementary Figure 4.**
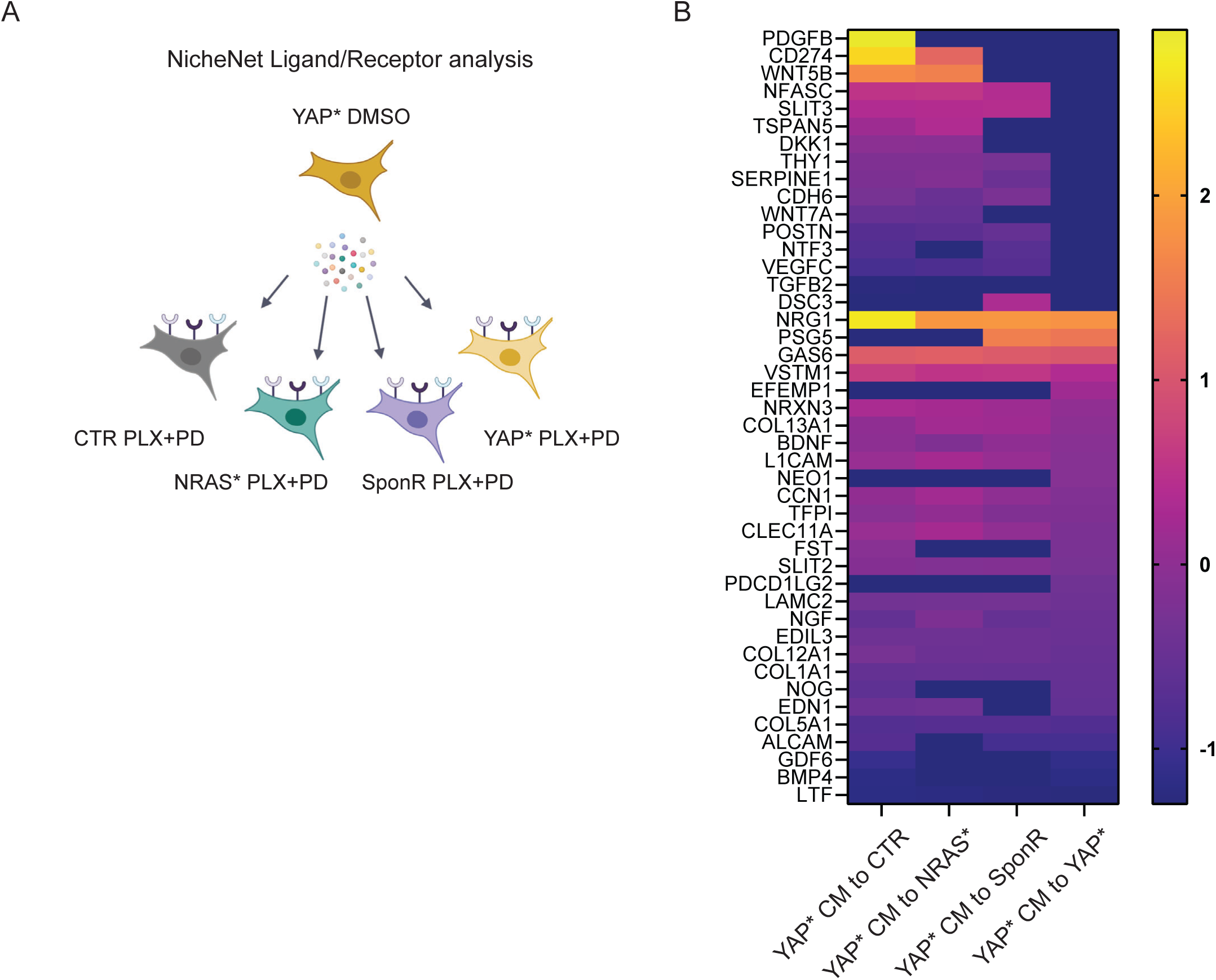
Co-cultures reveal inter-subclone crosstalk. A. Schematic illustration of the NicheNet analysis. Ligand expression by YAP* cells in DMSO was cross-referenced with receptor expression in CTR, NRAS*, YAP*, and SponR cells in PLX-PD. B. Heatmap of differential ligand expression in CTR, NRAS*, YAP*, and SponR cells inferred to be capable of driving gene expression signature linked to BRAFi + MEKi using NicheNet analysis

**Supplementary Figure 5.**
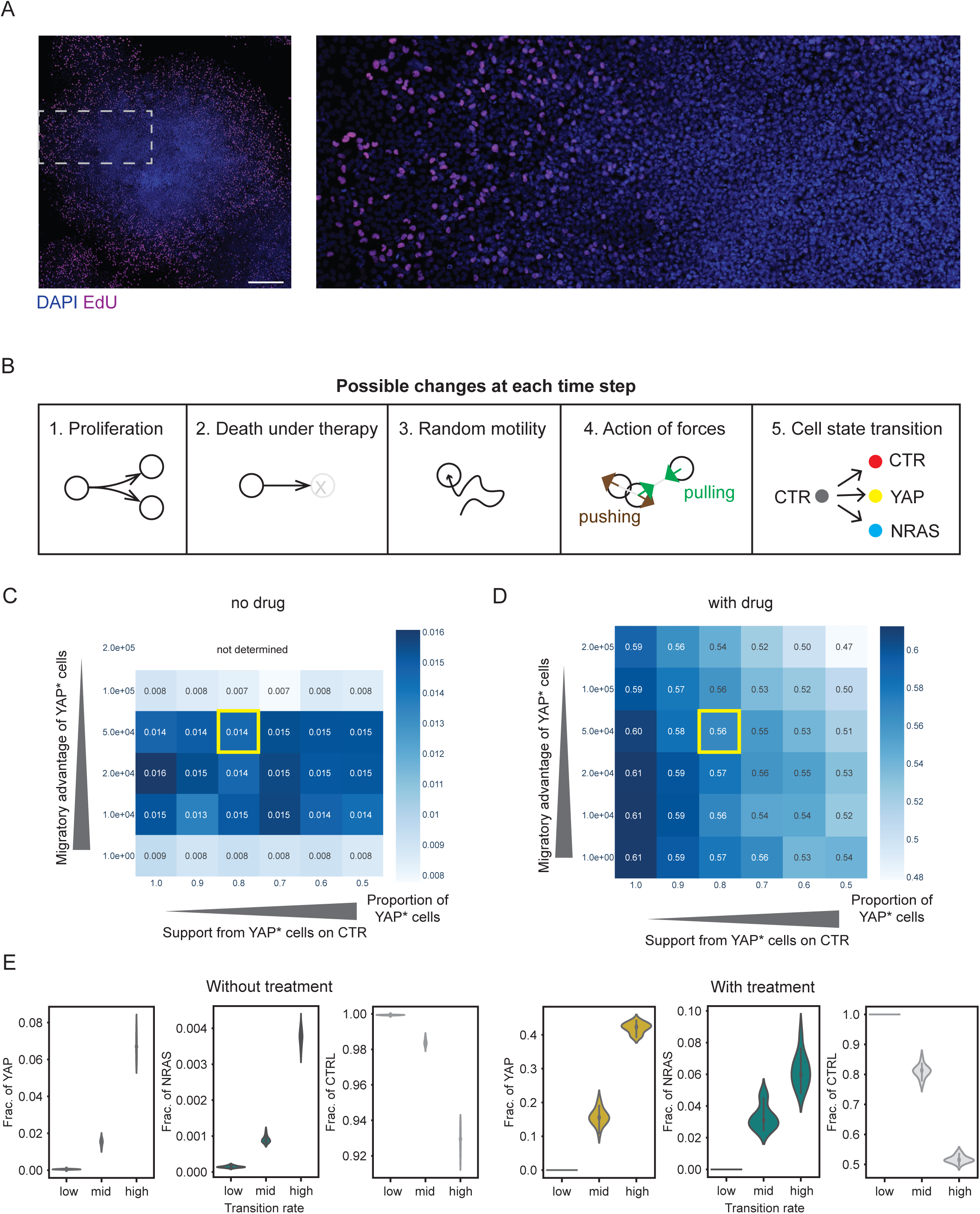
Computational modelling predicts tumour heterogeneity as favoured state following therapy. A. Images show EdU in magenta (proliferation) and DAPI in blue (DNA) in an A375 colony (left), with a zoomed view shown on the right. Scale bar is 500µm. B. Illustration showing the possible changes in cell state between time steps in the model. C. Heatmap shows the interplay of strength of YAP* self-inhibition (x-axis) and relative migratory advantage of YAP* cells (y-axis) in the absence of drug. Number shows the proportion of YAP* cells at the last time step. D. Heatmap shows the interplay of strength of YAP* self-inhibition (x-axis) and relative migratory advantage of YAP* cells (y-axis) in the presence of drug. Number shows the proportion of YAP* cells at the last time step. E. Plots show how varying the transition rate impacts the relative levels of YAP* and NRAS* cells in both the presence and absence of drug.

**Supplementary Figure 6.**
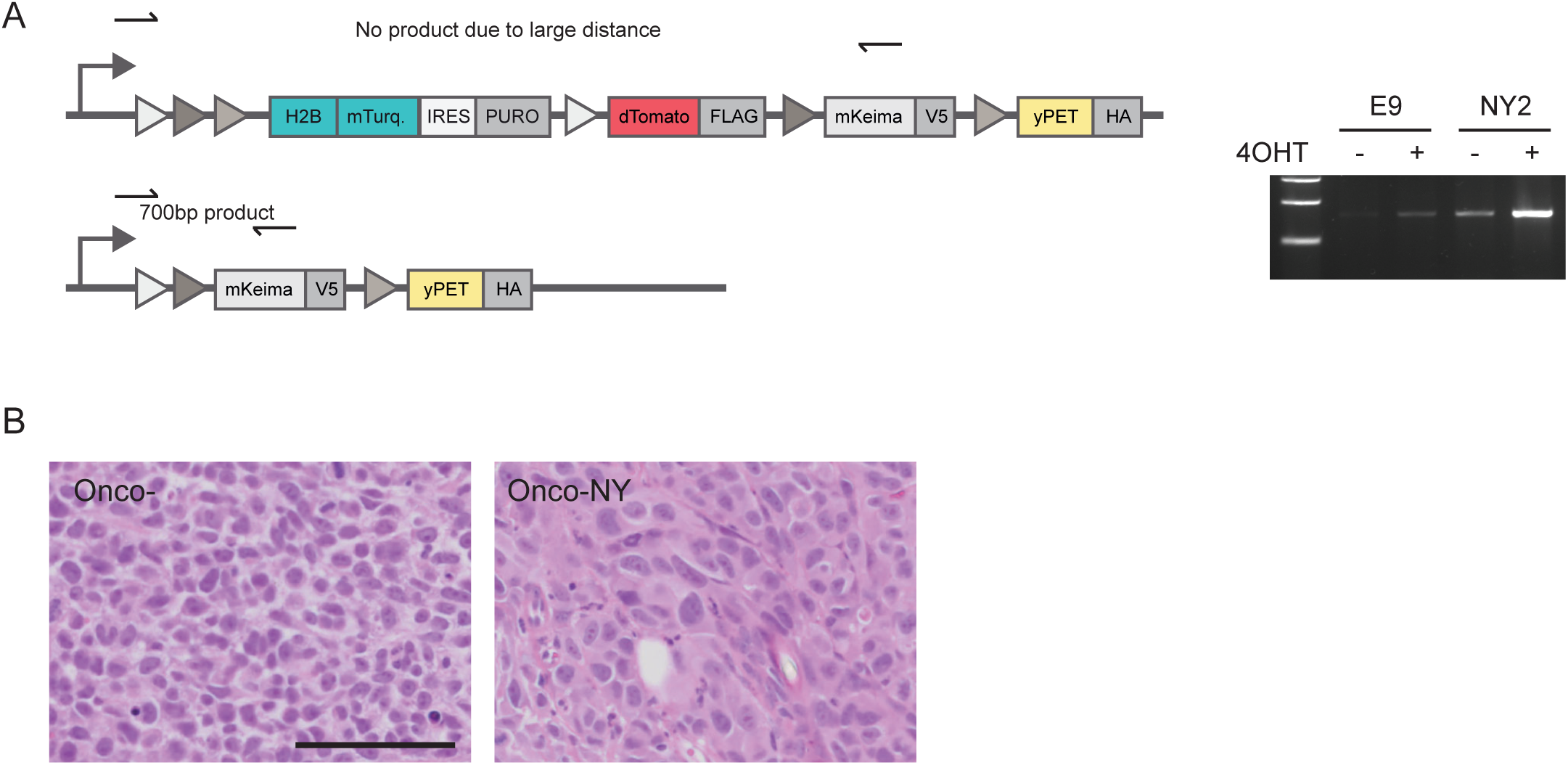
An experimental model of intra-tumour heterogeneity. A. Panel shows PCR strategy and PCR products generated from genomic DNA of Onco- and Onco-NY cells +/- 4OHT. B. H&E images of Onco-A375 and Onco-NY-A375 tumours

**Supplementary Figure 7.**
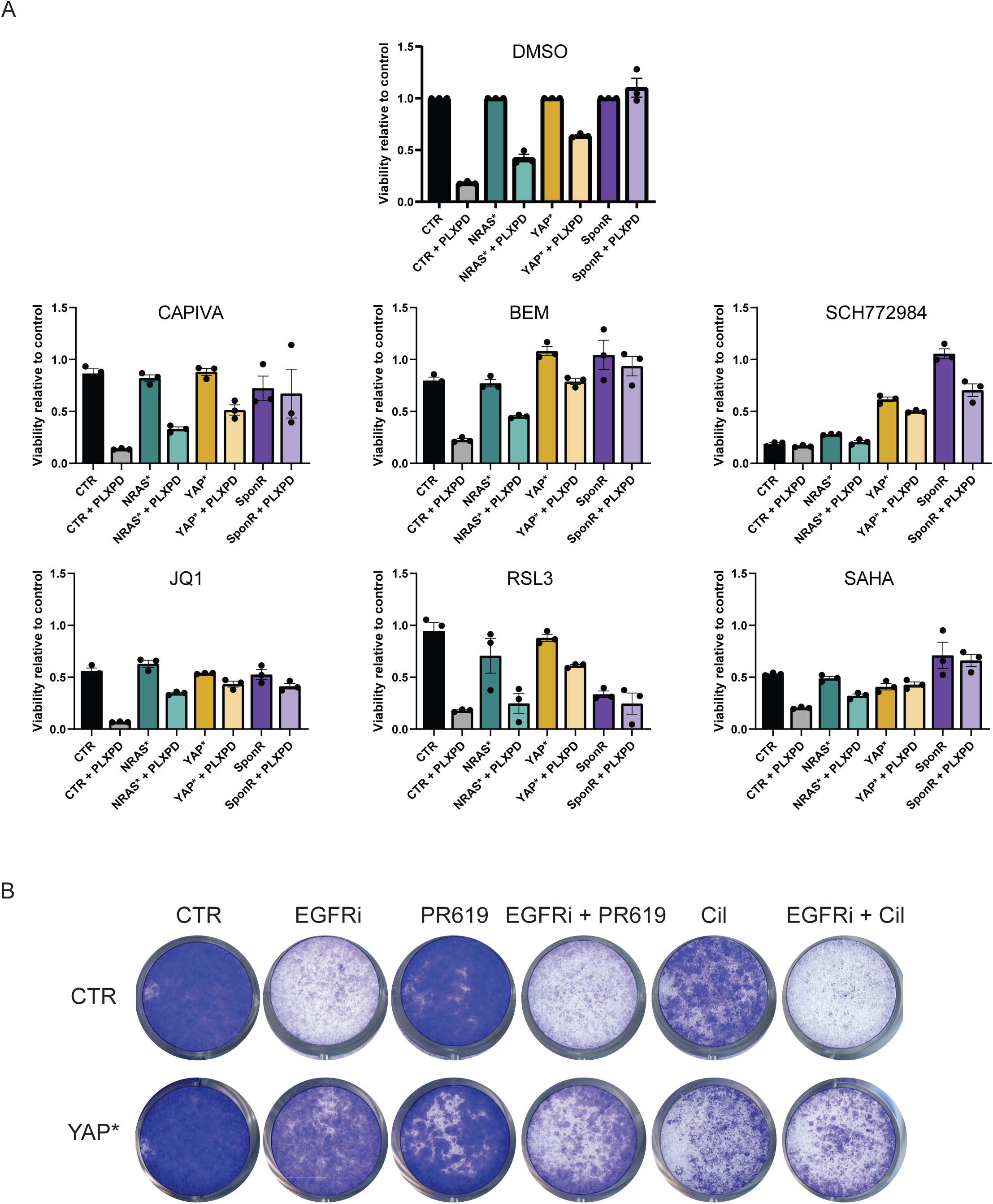
Identification vulnerabilities of different drug resistant states. A. Plots show the effect of compounds targeting ERK (SCH772984), HDACs – SAHA, Bromodomains – JQ1, AXL (Bemcetinib), GPX4 – RSL3, and AKT (Capivasertib) on CTR, NRAS, YAP*, and SponR cells with and without 1µM PLX4720 & 1µM PD184352. Plots are normalised to the number of CTR, NRAS*, YAP*, or SponR cells in the absence of any drug. The mean of three independent experiments is plotted. B. Images show crystal violet staining of CTR and YAP* H1975 treated for 8 days with the indicated inhibitors – EGFRi is 500nM Osimertinib and Cil is 10µM cilengitide.

## Notes

### Summary of Updates

We have added new data in Figure relating to heterogeneous cell states in human melanoma. Supplementary Movies are added. Formatting and typos have been addressed

